# Mll1 pioneers histone H3K4me3 deposition and promotes formation of CD8^+^ T stem cell memory

**DOI:** 10.1101/2023.01.18.524461

**Authors:** Adam J. Getzler, Megan A. Frederick, Justin J. Milner, Thomas Venables, Huitian Diao, Clara Toma, Shashank D. Nagaraja, Dominic S. Albao, Simon Bélanger, Shanel M. Tsuda, Jihye Kim, Shane Crotty, Ananda W. Goldrath, Matthew E. Pipkin

## Abstract

CD8^+^ T cells with stem cell-like properties (T_SCM_) sustain adaptive immunity to intracellular pathogens and tumors. However, the developmental origins and chromatin regulatory factors (CRFs) that establish their differentiation are unclear. Using an RNA interference screen of all CRFs we discovered the histone methylase Mll1 was required during T cell receptor (TCR) stimulation for development of a T_SCM_ precursor state and mature memory (T_MEM_) cells, but not short-lived or transitory effector cell-like states, in response to viral infections and tumors. Mll1 was essential for widespread *de novo* deposition of histone H3 lysine 4 trimethylation (H3K4me3) upon TCR stimulation, which accounted for 70% of all activation-induced sites in mature T_MEM_ cells. Mll1 promoted both H3K4me3 deposition and reduced TCR-induced Pol II pausing at genes whose single-cell transcriptional dynamics explained trajectories into nascent T_SCM_ precursor states during viral infection. Our results suggest Mll1-dependent control of Pol II elongation and H3K4me3 establishes and maintains differentiation of CD8^+^ T_SCM_ cell states.

## Introduction

Naive CD8 T cells activated by T cell receptor (TCR), co-receptor and cytokine stimulation during infections or cancer develop transcriptionally heterogeneous cell states (*1–5*) that give rise to diverse effector (T_EFF_), memory (T_MEM_), exhausted (T_EX_) and dysfunctional (T_DYS_) cell phylogenies (*6–10*). During acutely resolved infections in which antigen exposure is episodic, protective T_EFF_ and T_MEM_ cells develop (*8, 11*). Memory stem cells (T_SCM_) that develop after acute infection are an incompletely defined subset of central memory (T_CM_) T cells that highly express the transcription factor (TF) Tcf1 (encoded by *Tcf7*), re-express some naive cell attributes and sustain adaptive immunity by self-renewing under homeostatic conditions. In response to secondary infection, T_SCM_ cells proliferate most extensively and regenerate the compendium of protective T_EFF_ and T_MEM_ cell populations (*12–17*). These functions might explain the capacity of T_MEM_ cells to survive and proliferate to extents that exceed individual organismal lifespans (*18*). During chronic infections and cancer, subsets of “stem-like” progenitor T_SCM_-like cells (termed exhausted progenitor (T_EX_^prog^) cells) persist in the face of chronic antigen exposure and differentiate into transitory T_EFF_-like (T_EX_^trans^) cells that sustain the T cell response and curb pathogen or tumor loads (*19, 20*), but which progressively lose function and become terminally “exhausted” (T_EX_^term^) or “dysfunctional” (*21–27*). Differential chromatin remodeling correlates with the formation of distinct CD8 T cell states (*28–31*), but the transcriptional mechanics that create gene expression heterogeneity in TCR stimulated cells and establish their early developmental trajectories are incompletely understood. Moreover, while chromatin regulatory factors (CRFs) that silence genes which promote the longevity of T_MEM_ and that enforce terminal differentiation of T_EFF_ cells have been identified (*32–35*), those which establish transcriptional competence and promote formation of long-lived stem cell-like memory are still unknown.

## Results

### Identification of CRFs that coordinately regulate expression of PD-1, LAG3, TIM3 and Id3

Very early after either acute or chronic lymphocytic choriomeningitis virus (LCMV) infection, activated CD8 T cell populations that most highly express the transcriptional regulators Id3 and Tcf1 (encoded by *Tcf7*) have an enhanced capacity to develop into T_SCM_ or T_EX_^prog^ cells compared to T_EFF_ or T_EX_ cells (*2, 36*). In either infectious context, TCR activated cells initially uniformly induce cell surface expression of the co-inhibitory receptors PD-1 (*Pdcd1*), Lag3 (*Lag3*) and Tim3 (*Havcr2*) (*37*), but only those responding to chronic infection increase and sustain their expression (*2, 36, 37*). We hypothesized that CRFs which coordinately regulated expression of Id3, and the T_EX_ cell markers PD-1, Lag3 and Tim3 following initial TCR stimulation, might also more generally control the early formation of T_SCM_ and T_EX_^prog^ cells as opposed to more terminally differentiated T_EFF_ or T_EX_ cells. To screen for these CRFs, a library of retrovirally expressed microRNA30-embedded short-hairpin RNAs (shRNAmirs) targeting all murine CRFs was constructed and naive LCMV-specific (P14) CD8 T cells activated via TCR signals were transduced with individual shRNAmirs in separate wells (Fig 1A). The expression of nine cell surface receptors and a GFP reporter allele in the endogenous *Id3* locus (Id3-GFP) was monitored in shRNAmir-transduced cells, and the average loss-of-function effects on expression of these T cell differentiation markers derived from multiple, unique shRNAmirs targeting each CRF in the separate wells were summarized (RNAi-effects) and ranked (fig S1, table S1). We performed two orthogonal screens: The first examined surface receptor expression on P14 cells 6 days after primary TCR-activation and culture with interleukin-2 (IL-2) under conditions that promote key aspects of differentiation into effector (IL-2^hi^) or memory (IL-2^lo^) cells (Fig 1B *right*, fig S1A-B, table S1)(*38*); the second analyzed P14 *Id3*-GFP CD8 T cells 3 days after primary TCR stimulation (two days after transduction) and focused on expression of *Id3*-GFP, PD-1 and Lag3 (Fig 1B *left*, fig S1A-B, table S1).

**Fig 1.**
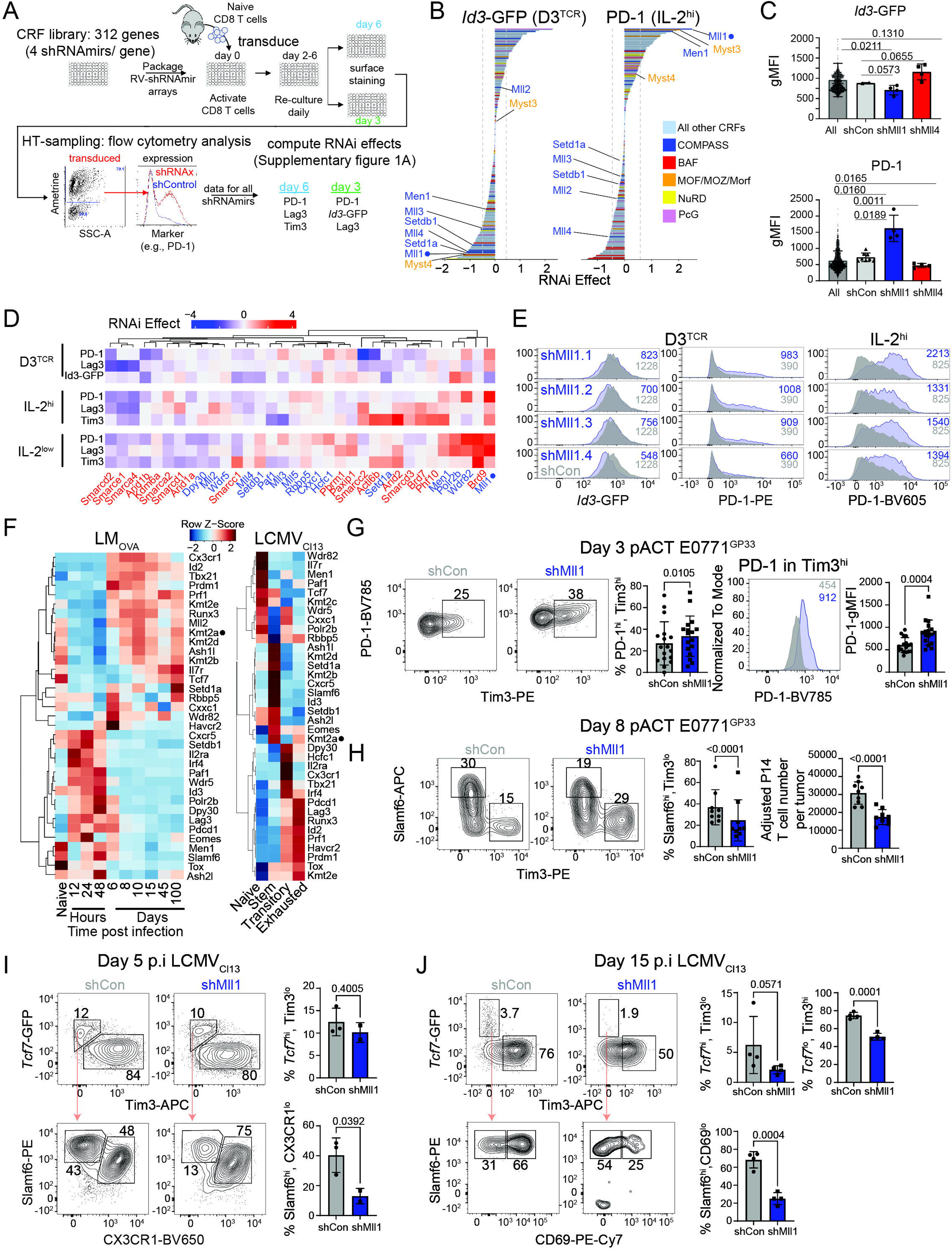
**Identification of Mll1 as an essential regulator of progenitor T_SCM_-like cell formation** (**A**) Arrayed RNA interference screening layout. (**B**) The average RNAi effect resulting from suppression of individual CRFs on expression of *Id3*-GFP (left, d3^TCR^) and PD-1 (right, IL-2^hi^) in each screen. Dashed line indicates top and bottom 15^th^ percentile. (**C**) Bar charts show the mean raw expression values (geometric mean fluorescent intensity) of *Id3*-GFP (top) and PD-1 (bottom) in response to each cognate shRNAmirs (symbols). P values were calculated using Welch’s t-test. (**D**) RNAi-effects for COMPASS (blue) and BAF (red) complex subunits in each screen were clustered by Euclidean distance. (**E**) Representative flow cytometry data of individual shRNAmirs targeting a non-expressed gene (shCon, gray) or Mll1 (shMll1, blue) in each screening condition. Numbers in plots indicate geometric mean fluorescence intensity. (**F**) Heatmaps show normalized transcript counts of selected COMPASS, COMPASS-like, T_SCM_ and T_EX_ cell signature genes in purified antigen-specific CD8 T cells responding to Listeria monocytogenes-Ova (LM-Ova) (*45*), or LCMV_Cl13_ (*25*). Intensity of color represents highest expression of an individual gene across columns. (**G-H**) Representative flow cytometry plots after gating on P14 CD8^+^ T cells transduced with shRNAmirs specific for negative control genes (shCon) or Mll1 (shMll1) in E0771^GP33^ tumors, and comprise concatenated data from multiple mice (n=3). Histogram overlay shows geometric mean fluorescent intensity (gMFI); gating is indicated. Bar charts summarize the means of individual mice (symbols). P values calculated via paired Student’s t-test. (**I-J**) Representative flow cytometry plots after gating on adoptively co-transferred P14*^Tcf7^*^-GFP^ CD8^+^ T cells transduced with either control (shCon) or Mll1 (shMll1) specific shRNAmirs from the spleens of host mice 5 or 15 days after LCMV_Cl13_ infection. Data was concatenated from multiple recipient mice (I, n=3; J, n=4). Tcf7^hi^, Tim3^lo^ cells (top) were gated and analyzed for Slamf6 and CD69 expression (bottom). Bar charts summarize the means of individual mice (symbols). P values calculated via Welch’s t-test, except for J (top), which used the Mann-Whitney test.

Multiple genes encoding CRF subunits that assemble into the same biochemical complexes exhibited analogous RNAi-effects, such that suppressing each CRF either increased (positive RNAi-effect) or decreased (negative RNAi-effect) expression of PD-1, Lag3, Tim3 or *Id3*-GFP (fig S1C, table S1). Genes encoding the COMPASS-family of histone methyltransferase complexes (*39*), MOZ/MOF acetyltransferases, NuRD and polycomb group (PcG) proteins were significantly enriched among all CRFs with positive PD-1 RNAi-effects, whereas genes encoding BAF complex subunits were enriched among CRFs with negative PD-1 RNAi-effects (Fig 1B, fig S1B-C). In addition, CRFs with positive PD-1 RNAi-effects positively correlated with those that exhibited positive Lag3 or Tim3 RNAi-effects, and modestly correlated with those that had negative RNAi-effects on *Id3*-GFP (fig S1D). Thus, multiple distinct CRFs that assemble in shared complexes regulated the coordinated, and reciprocal, expression of co-inhibitory receptors and *Id3*-GFP upon naive CD8 T cell activation.

Individual CRFs were prioritized by focusing on those whose depletion had both the strongest effects and coordinately increased or decreased expression of more than one co-inhibitory receptor (Fig 1B and fig S1B, top/bottom 15% of RNAi-effects, table S1). Depletion of Mll1, the catalytic subunit of Mll-COMPASS-like complexes (encoded by *Kmt2a*), increased expression of PD-1 and Lag3 in IL-2^lo^ cultures, as well as, PD-1, Lag3 and Tim3 in IL-2_hi_ cultures (Fig 1B-E and fig S1B, S2A, table S1). Depletion of Menin (*Men1*), a key component of Mll1 and Mll2 (*Kmt2b*) -COMPASS-like complexes (*40, 41*) exhibited similar effects (Fig 1B and fig S1B). In addition, suppression of the catalytic acetyltransferase MOZ (Myst3/ *Kat6a*), which associates with Mll1-COMPASS-like complexes, and its paralog MORF (Myst4/*Kat6b*) (*42*) also increased both PD-1 and Lag3 expression (Fig 1B and fig S1B, table S1). However, depletion of Mll2, the closest paralog of Mll1 (*39*), exhibited distinct effects (Fig 1B-D, fig S1B and fig S2A, table S1), which implies that Mll1 and Mll2 each have specific functions in CD8 T cells, analogous to their distinct requirements in other cell types (*43*). Transduction of single-guide RNAs (sgRNAs) specific for Mll1/*Kmt2a* into P14 Cas9-transgenic CD8 T cells also caused upregulation of PD-1 and Tim3 (fig S2B). Immunoblotting confirmed the on-target activity of all *Kmt2a*-specific shRNAmirs and sgRNA/Cas9-mediated disruption of *Kmt2a* (fig S3A). Thus, an Mll1-specific COMPASS-like complex might be required for controlling co-inhibitory receptor expression during TCR activation.

In the second screen, Mll1 RNAi-effects were also in the top 15% of CRFs whose depletion caused upregulation of both PD-1 and Lag3, and that impaired Id3-GFP expression (Fig 1B and fig S1B). In this setting, depletion of Mll3 (*Kmt2c*), Mll4 (*Kmt2d*), Setd1a (*Setd1a*), or Setdb1 (*Setdb1*) also impaired upregulation of *Id3*-GFP (Fig 1B, 1D, and fig S1B, table S1), suggesting that multiple Mll-COMPASS-like or Set1-COMPASS complex activities might concertedly or redundantly promote *Id3* expression during initial TCR stimulation. Additionally, depletion of Mll1 in P14 cells encoding GFP in the *Tcf7* locus (P14*^Tcf7^*^-GFP^) impaired *Tcf7*-GFP expression compared to control-transduced cells (fig S3B). As expression of *Tcf7* correlates with and promotes development of cells that express Id3 (*36, 44*), the impaired Id3 expression in the absence of Mll1 could involve reduced Tcf1 expression. Notably, Mll1-deficient cells were not generically defective, because depletion of Mll1 did not impair cell division history or overall accumulation during the first week of cell culture, or the ability to express key cytokines that are induced by T_EFF_ and T_MEM_ cells upon TCR-like re-stimulation (fig S3C-D). Thus, Mll1 is required during TCR stimulation of naive CD8 T cells *in vitro* to activate Id3 expression and to restrain expression of PD-1, Lag3 and Tim3.

Consistent with the phenotypic effects of Mll1-deficiency after TCR stimulation *in vitro*, analysis of previously published *ex vivo* gene expression data indicated *Kmt2a* expression was highest in naive cells and after day 8 post infection (pi) and in long-lived T_MEM_ cells following intracellular bacterial infection (*45*), and clustered with expression of the T_MEM_- and T_SCM_-associated genes *Tcf7* and *Il7r* (Fig 1F, *left*). *Kmt2a* expression was also highest in stem-like CD8 T cells after chronic LCMV infection (*19*), and clustered with expression of *Tcf7*, *Id3, Slamf6* and *Cxcr5* (Fig 1F, *right*). Conversely, *Kmt2a* was less highly expressed in T_EFF_ cells isolated from the early phase of acute infection (Fig 1F, *left*), and in both T_EFF_^trans^ and

T_EX_^term^ cells during chronic infection, and clustered apart from *Pdcd1* (PD-1), *Havcr2* (Tim3) and *Lag3* (Lag3), which were highly expressed in these cells (Fig 1F, *right*). Thus, *Kmt2a* expression is dynamic and positively correlates with formation of the most long-lived, protective T cell subsets that develop during both acute and chronic infection.

### Mll1 is essential for development of T cell states that sustain responses in tumors and during chronic viral infection

To define *in vivo* requirements of Mll1, the effects of its depletion in CD8 T cells responding to engineered solid tumors expressing the GP_33-41_ epitope of LCMV (GP33) were examined in mice. P14 TCR transgenic CD8 T cells recognize GP33 and were transduced with shRNAmirs specific for Mll1 (shMll1) or negative control mRNAs (*Cd4* or *Cd19*) that are not expressed in CD8 T cells (shCon), and were co-transferred into wild type mice engrafted with B16 melanoma or E0771 triple negative breast tumors expressing GP33 fused to GFP (B16^GP33^ or E0771^GP33^), and their parental controls expressing GFP-only (B16^GFP^ or E0771^GFP^) (fig S4A). In these models, P14 cells accumulated systemically post adoptive cell transfer (pACT), but preferentially expanded and upregulated PD-1 in GP33-GFP expressing tumors, compared to the spleen or parental GFP-expressing tumors (fig S4B and fig S4C). Thus, the phenotypes of transferred P14 cells *in vivo* were induced specifically by antigen display in the tumor. Compared to shCon-transduced P14 cells in E0771^GP33^ and B16^GP33^ tumors, shMll1-transduced P14 cells formed significantly greater fractions of PD-1^hi^ Tim3^hi^ cells (Fig 1G, p=0.0105; fig S3D, p=0.0081), and those responding to E0771^GP33^ tumors exhibited higher PD-1 expression among gated Tim3^hi^ cells (Fig 1G p=0.0004, fig S4E), and initially accumulated as well as, or better, than shCon-transduced cells by day 3 pACT (fig S4F). Notably, shMll1-transduced P14 cells initially delayed E0771^GP33+^ tumor growth compared to transfer of shCon-transduced P14 cells, but the tumors ultimately regressed analogous to recipients of shCon-transduced P14 cells (fig S4G). Thus, Mll1-deficient cells preferentially adopted T_EFF_ cell-like states and were transiently functional in the tumor microenvironment, indicating they were not generically defective, but could not sustain tumor control.

Tumor-specific CD8 T cells that are Tcf1^hi^ Slamf6^hi^ and Tim3^lo^ normally define a progenitor cell population with T_SCM_-like properties that sustains the T cell response in tumors (*27*). However, shMll1-transduced P14^Tcf7-GFP^ donor cells in E0771^GP33+^ tumors expressed less *Tcf7*-GFP overall on day 3 pACT (fig S4H, p=0.0732), even those that were Tim3^lo^ (fig S4H, p=0.0212), compared to control-transduced P14*^Tcf7^*^-GFP^ cells. By day 8 pACT, shMll1-transduced cells generated significantly smaller fractions and numbers of Slamf6^hi^ Tim3^lo^ cells, compared to shCon-transduced cells (Fig 1H, p<0.0001). These results indicate that Mll1 is necessary for the early formation and maintenance of progenitor T_SCM_-like cells and the persistence of responding CD8 T cells in the tumor microenvironment.

The tumor microenvironment could differentially affect the development of Mll1-deficient cells. Therefore, Mll1-depleted P14 cells were analyzed after transfer into wildtype hosts followed by infection with LCMV Clone 13 (LCMV_Cl13_), which causes chronic viremia (*46*). Very early after infection, activated cells develop into Tcf7^hi^ Id3^hi^ Tim3^lo^ exhausted cell precursor and Tcf7^lo^ Id3^lo^ Tim3^hi^ effector cell precursor populations (*36*), that ultimately establish a continuum of cell states termed T_EX_^prog^, transitory effector/exhausted intermediate (T_EX_^trans^/T_EX_^int^) and terminally exhausted (T_EX_^term^) cells as the infection becomes chronic (*24–26*). The CRFs that initiate this pattern early during infection are unknown. On day 5 post infection (pi), both shCon-transduced and shMll1-transduced P14 cells equivalently polarized into populations of *Tcf7*-GFP^hi^ Tim3^lo^ (T_EX_-precursor) and *Tcf7*-GFP^lo^ Tim3^hi^ (T_EFF_-precursor) cells (Fig 1I, *top*). Notably, shCon-transduced T_EX_ precursors were comprised of equivalent fractions of Slamf6^hi^ CX3CR1^lo^ and Slamf6^hi^ CX3CR1^hi^ cells (Fig 1I, *top*), suggesting T_EX_ precursors manifested roughly equal bias toward future T_EX_^prog^ (CX3CR1^lo^) and T_EX_^trans^/T_EX_^int^ (CX3CR1^hi^) states at this time. In contrast, shMll1-transduced *Tcf7*-GFP^hi^ Tim3^lo^ cells were predominantly Slamf6^int/lo^ CX3CR1^hi^, indicating Mll1-deficiency skewed development toward the T_EX_^trans^/T_EX_^int^ state (Fig 1I, *bottom* p=0.0392). This suggests Mll1 is necessary to initially form or stabilize precursors that typically develop into T_EX_^prog^-like cells. Supporting this conclusion, by day 15 pi, the overall number and fraction of shMll1-transduced P14 cells among total CD8^+^ cells accumulated less and were not sustained compared to shCon-transduced cells (fig S5A). In addition, shMll1-transduced cells formed reduced fractions of *Tcf7*-GFP^hi^ Tim3^lo^ cells compared to shCon-transduced cells (Fig 1J, *top*, p=0.0571), and fewer shMll1-transduced cells that were *Tcf7*-GFP^hi^ were also CD69^hi^ compared to shCon-transduced cells at this time (Fig 1J, *bottom* p=0.0004). Thus, Mll1-deficient cells form fewer *Tcf7*-GFP^hi^ cells that develop a phenotype corresponding to *bona fide* T_EX_^prog1^ cells (*26*).

Although Mll1-deficient cells expressed much less Slamf6 overall on day 15 pi (fig S5B), suggesting they were more effector-like, shMll-transduced cells formed smaller fractions of *Tcf7*-GFP^lo^ Tim3^hi^ cells compared to shCon-transduced cells (Fig 1J, p = 0.0001), and these were uniformly CX3CR1^hi^ (fig S5C, p=0.0230), which is atypical of canonical T_EX_^trans^-like cells (*26*). In addition, whereas most Slamf6^int/lo^ shCon-transduced cells were CD69^hi^, a characteristic of T_EX_^term^ cells (*26*), most Slamf6^int/lo^ shMll1-transduced cells were CD69^lo^ (fig S5B, p =0.0052). These results indicate that cells lacking Mll1 develop into non-canonical effector-like states, which do not efficiently mature into the fully exhausted phenotype during chronic infection.

### Mll1 is essential for establishing the normal pattern of circulating T_MEM_ cells during acute viral infection

Strong or persistent TCR stimulation increases development of T_EX_ precursors during chronic infection (*36*) and could exacerbate or modify defects in Mll1-deficient cells. To examine if Mll1 was important for the formation of protective T_MEM_ cells, which normally develop when infections resolve rapidly and antigen is completely cleared (*11, 47*), naive shCon-transduced and shMll1-transduced P14 CD8 T cells were co-transferred to wildtype host mice subjected to LCMV Armstrong (LCMV_Arm_) infection, which acutely resolves (*46*). In this setting, Mll1-deficient cells formed 2-4x fewer T_EFF_ cells at the normal peak of the response compared to control cells (Fig 2A), but subsequently decreased more sharply than control cells afterward, and formed ∼10-fold fewer total T_MEM_ cells (Fig 2A).

**Fig. 2.**
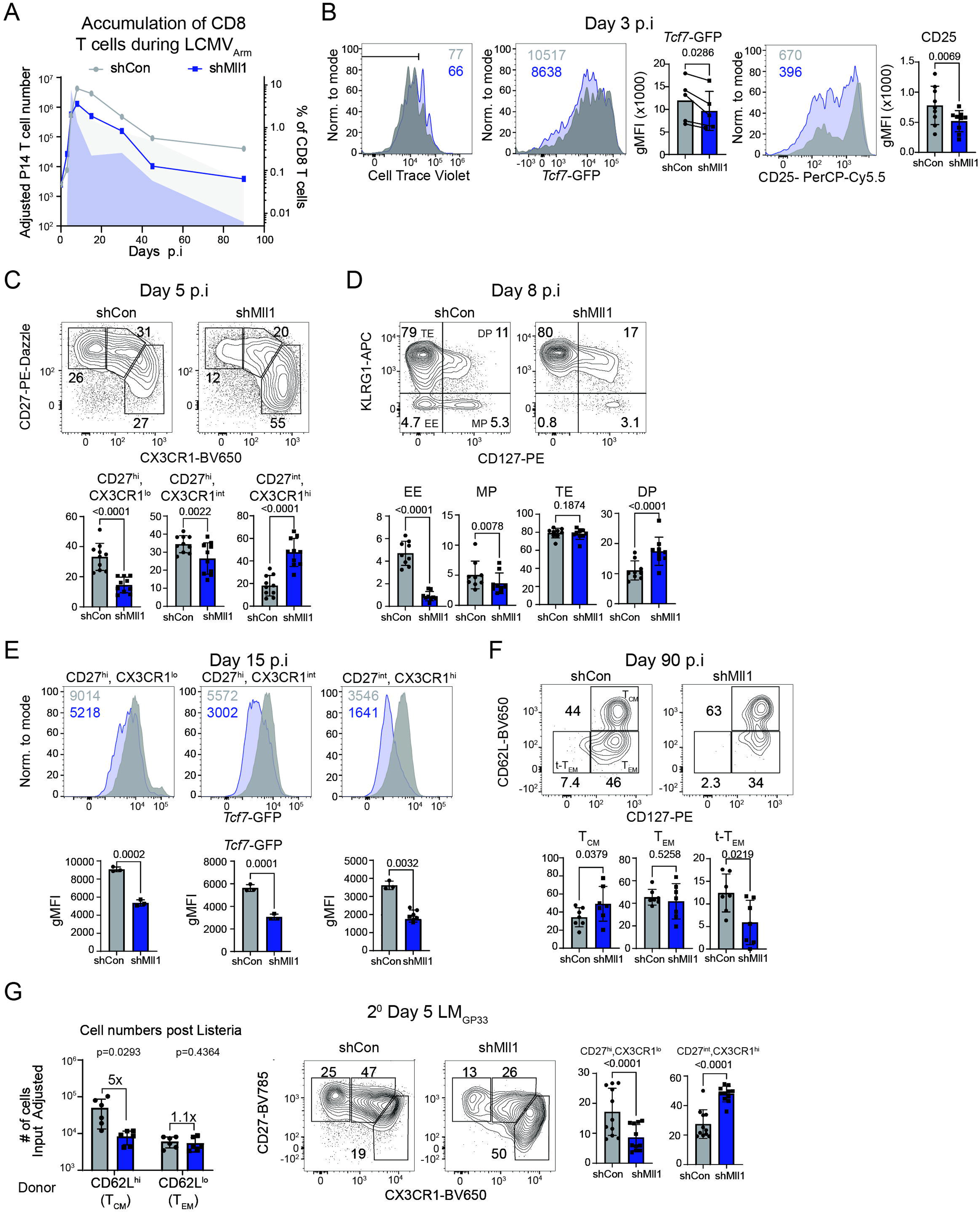
**Mll1 is essential for development of T_MEM_ cells following acute viral infection.** P14 or P14*^Tcf7^*^-GFP^ CD8^+^ T cells transduced with either control (shCon) or Mll1 (shMll1) specific shRNAmirs were adoptively co-transferred into wildtype host mice subjected to LCMV_Arm_ infection (A-F). (**A**) P14 cell accumulation. Symbols indicate normalized transduced P14 cell numbers in the spleen. Shaded plot indicates normalized percent of transduced P14 cells of all CD8^+^ T cells. Both data present the mean of n≥3 individual mice. (**B**) Naive P14 cells were cell trace violet (CTV) labeled before transduction, and the percent of each transduced population that underwent more than the median number of divisions by day 3 pi in the spleen is indicated in the plot (left). Histograms show the gMFI of *Tcf7*-GFP (middle) and CD25 expression (right). Numbers in plots indicate gMFI of each marker, which are summarized in bar charts as the means for all mice (symbols). Paired t-test used to compare means. Representative flow cytometry data concatenated from four recipient mice. (**C-F**) Representative flow cytometry of transduced P14 CD8^+^ T cells in the spleen are data concatenated from 3-5 mice. Values in plots indicate frequencies, and are summarized as means in bar charts for all mice (symbols). Mice with less than 10 transduced cells (day 90 pi) were excluded from statistical analyses of cell frequencies, and the mean normalized transduced P14 cell numbers per spleen were summarized for individual mice (symbols) in (F). Paired t-test or Wilcoxon test were used to compare means. (**E**) The representative histograms analyzed expression *Tcf7*-GFP in phenotypic populations after gating on transduced P14*^Tcf7^*^-GFP^ CD8^+^ T cells. Numbers in plots indicate gMFI and are summarized for all mice (symbols). Paired t-test used for statistics. (**G**) The accumulation of donor derived CD62L^hi^ or CD62L^lo^ memory cells from day 90 pi that were either shCon or shMll1 transduced were mixed, adoptively transferred to naive wildtype hosts and subjected to LM^GP33^ infection. The abundance (left) and phenotype (right) of transduced P14 cells were analyzed on 5 pi in the spleen. Transduced P14 cell numbers and frequencies of phenotypic subsets were summarized for all mice (symbols) in bar charts. Paired t-tests used for statistics.

The T_EFF_ cell population becomes heterogeneous before acute infections resolve, and only some T_EFF_ cells develop into precursors of T_MEM_ cells (*3, 12, 36, 48-51*), but how this heterogeneity is initially established is still unclear. Analysis of P14 cells labelled with cell trace violet (CTV) showed that Mll1-deficient cells initially accumulated but underwent fewer cell divisions, less efficiently sustained high *Tcf7*-GFP expression (Fig 2B *left/middle*, p =0.0286) and expressed less CD25 compared to control-transduced cells on day 3 pi (Fig 2B *right*, p= 0.0069). This indicated that the typical inverse relationship between CD25 and Tcf7 expression did not polarize in the absence of Mll1 (*16*). Furthermore, compared to shCon-transduced P14 cells on days 5 and 15 pi, those transduced with shMll1 developed reduced fractions of CD27^hi^ CX3CR1^lo/int^ and CD27^int^ CX3CR1^int^ cells, which normally harbor precursors of central memory (T_CM_) cells (Fig 2C, p<0.0001,p=0.0022) (*52*), were mainly KLRG1^hi^ (Fig S6A, p<0.0001) and comprised a much larger fraction of CD27^lo^ CX3CR1^hi^ cells (Fig 2C, p<0.0001), which normally comprise precursors of T_EM_ cells and terminally differentiated effector (TE) cells (*51–53*). Although shMll1-transduced cells were as abundant as shCon-transduced P14 cells in the spleen on day 5 pi (Fig 2A), they incorporated less EdU, indicating they were proliferating less at this time (Fig S6B, top, p=0.0004). By day 8 pi, shMll1-transduced P14 cells formed lower frequencies of KLRG1^lo^ CD127^hi^ classical memory precursor (MP) cells (*51*) compared to shCon-transduced cells, and were virtually devoid of KLRG1^lo^ CD127^lo^ “early effector” (EE) cells (Fig 2D, p=0.0078, p<0.0001). These phenotypes were confirmed using P14-Cas9 transgenic cells transduced with Mll1-specific single-guide RNAs (sgMll1, fig S6C, p=0.2507, p=0.0216). In addition, KLRG1^hi^ shMll1-transduced P14 cells incorporated less EdU on day 8 pi (Fig S6D, p=0.0146), indicating they were less proliferative than shCon transduced KLRG1^hi^ cells at the peak of the response. Altogether these results suggest Mll1-deficiency prevented initial formation of conventional precursors of recirculating T_CM_ and T_EM_ cells, and promoted terminal differentiation.

T_EFF_ cells that most highly express *Tcf7* at the peak response to acute viral infection appear to be T_SCM_ precursors that form T_CM_ cells (*12*). On day 15 pi, all phenotypic subsets of shMll1-transduced P14^Tcf7-GFP^ cells expressed significantly less *Tcf7*-GFP compared to shCon-transduced cells (Fig 2E). Over time, the frequencies of shMll1-transduced P14 cells exhibiting a terminal T_EM_ (t-T_EM_) phenotype (CD62L^lo^ CD127^lo^) decreased (Fig 2F, *right*), those exhibiting a T_EM_ (CD62L^lo^ CD127^hi^) phenotype were mainly constant (Fig 2F, *middle*), and those with a central memory (T_CM_) phenotype (CD62L^hi^ CD127^hi^) increased (Fig 2F, *left*). However, the numbers of each phenotypic subset decreased strongly (fig S6E), as might be expected from their reduced *Tcf7*-GFP expression (*17*). We considered that the preferential persistence of shMll1-transduced cells with a T_CM_-phenotype was because CD27^hi^ T_EFF_ cells retained some *Tcf7*-GFP expression, suggesting they could have developed into functional T_CM_ cells. To test this, CD62L^hi^ and CD62L^lo^ shCon-transduced and shMll1-transduced P14 cells were isolated 90 days after primary LCMV_Arm_ infection, and 1:1 mixtures of shCon-transduced and shMll1-transduced cells that were either CD62L^hi^ or CD62L^lo^ were co-transferred to new naive hosts, and were then infected with *Listeria monocytogenes*-GP33 (LM_GP33_) to mimic a secondary infection (Fig 2G and fig S6F). On day 5 after LM_GP33_ infection, CD62L^hi^ donor P14 cells that were shMll1-transduced accumulated 5-10 fold less than shCon-transduced cells and formed a reduced fraction of secondary CD27^hi^ CX3CR1^lo^ T_CM_ precursors (Fig 2G p<0.0001). In contrast, the accumulation of both shMll1-transduced and shCon-transduced CD62L^lo^ donor cells was similar, but much less than the CD62L^hi^ transferred cells (Fig 2G and fig S6F), as expected of T_EM_ cells (*54*). Thus, Mll1 is essential for establishing Tcf7^hi^ memory precursors during the effector response and for forming *bona fide* CD62L^hi^ T_CM_ cells in the memory phase.

### Mll1 coordinately induces T_MEM_ gene expression and represses TE gene expression

To examine how Mll1 controls gene expression before T_CM_ cells definitively form, RNA-seq was used to analyze shMll1-transduced or shCon-transduced P14 T_EFF_ cells in the spleen that were either KLRG1^lo^ or KLRG1^hi^ 5 days after LCMV_Arm_ infection (fig S7A, and table S2). Analysis of shCon-transduced cells indicated KLRG1^lo^ cells more strongly expressed signatures characteristic of stem or progenitor stem-like cells, whereas KLRG1^hi^ cells more strongly expressed signatures of day 8 KLRG1^lo^ CD127^lo^ early effector (EE) and TE cells (fig S7B). TCR-induced gene expression (Fig 3A, *TCR dependent*) and the *Tcf7*^hi^ effector cell signature (*12*) were less strongly upregulated by shMll1-transduced cells compared to shCon-transduced cells in both KLRG1^lo^ and KLRG1^hi^ populations (Fig 3A, *Day 8 LCMV_Arm_*). The genes *Bcl6, Id3, Jund* and *Il6ra* in KLRG1^lo^ cells are normally more highly expressed in MP cells then TE(*55*), but were less highly expressed in shMll1-transduced KLRG1^lo^ cells compared to shCon-transduced (fig S7C, *left*, table S2). In addition, gene expression driven by Tcf1 and Tox, which is positively associated with progenitor T_SCM_-like cells, was less highly expressed in shMll1-transduced KLRG1^hi^ cells, compared to shCon-transduced cells (Fig 3A, *TCR dependent*). Reciprocally, shMll1-transduced cells overexpressed the signature of TE cells in both KLRG1^lo^ and KLRG1^hi^ populations (Fig 3A, *Day 8 LCMV_Arm_*), including the TE-signature genes *Zeb2, Bhlhe40*, *Ccl5* and *Cx3cr1*, compared to shCon-transduced cells and they also reactivated genes that are normally expressed in naive cells (Fig 3A, *TCR dependent*, fig S7C, *right* and table S2). Thus, Mll1 is necessary to activate expression of factors in both KLRG1^lo^ and KLRG1^hi^ cells that normally promote formation of long-lived T_MEM_ cells, and to repress aberrant co-expression of genes expressed by naive cells and TE cells.

**Fig 3:**
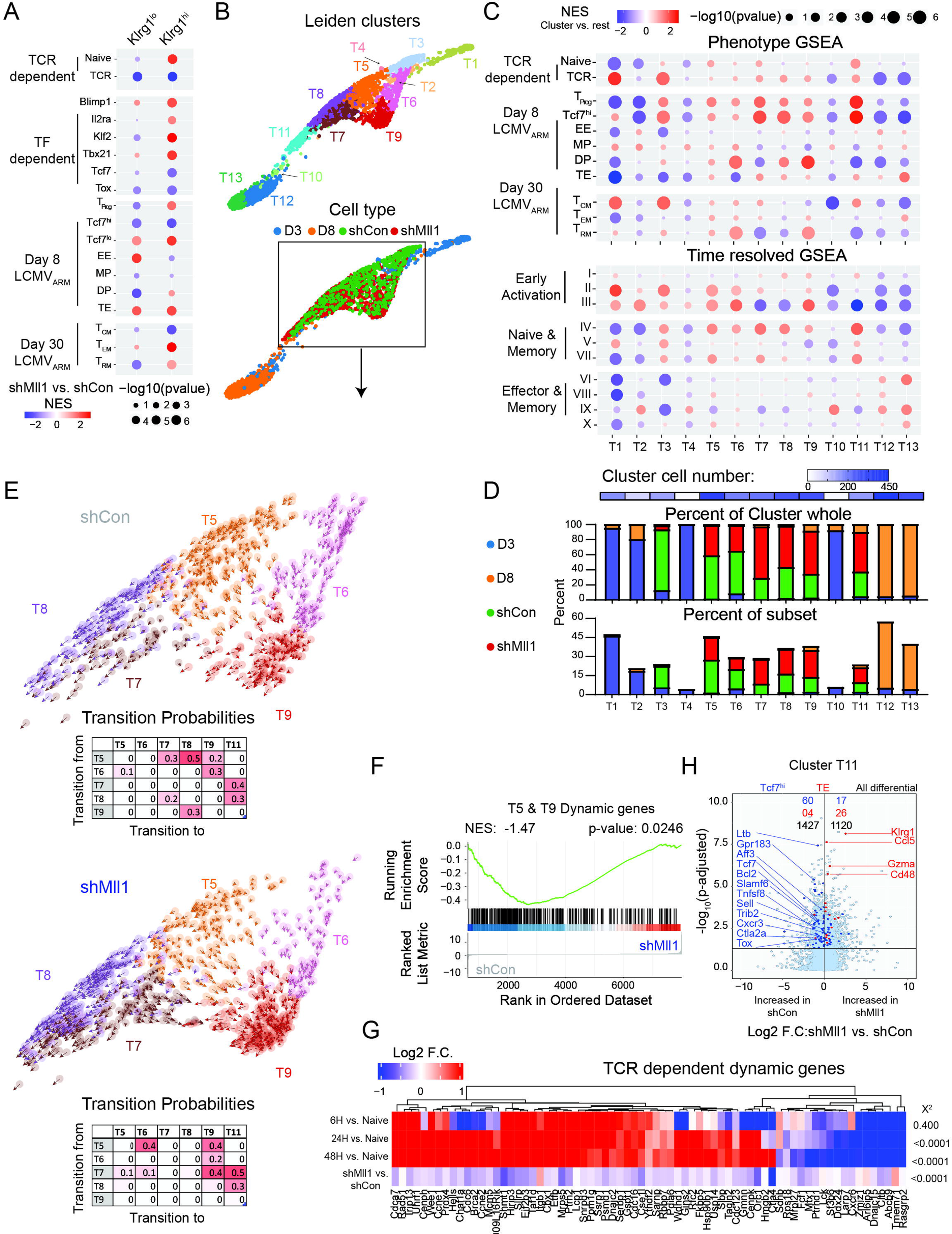
**Mll1 promotes transcriptional dynamics that establish a nascent T_SCM_ precursor state** (**A**) RNA-seq analysis examined shCon and shMll1-transduced KLRG1^lo^ and KLRG1^hi^ P14 cells in the spleen 5 days after LCMV_Arm_ infection. The enrichment of specific gene signatures (left) were analyzed against the Wald statistic of shMll1 vs shCon-transduced cells. (**B**) PAGA initialized force assisted analysis of the scRNA-seq data depicts Leiden clusters (top) and the cells of origin (bottom). (**C**) The specific expression of gene signatures (indicated on left) (*1, 2, 12, 45, 55, 68*) were analyzed within each Leiden cluster using Wilcoxon ranked score of the change in expression within each cluster versus all other clusters. (**D**) Cell numbers in each Leiden cluster are indicated (blue heatmap, top). Stacked bar charts indicate the frequency distribution of cells from each sample in Leiden clusters (middle), and the fraction of cells from each sample in each cluster (bottom). (**E**) RNA velocity vector fields for shCon (top) and shMll1 (bottom)-transduced cells in clusters T5-T9. The table inset indicates the intercluster transition potentials. Note distinct transition potentials from cluster T5 in shCon versus shMll1-transduced cells. (**F**) The bias in expression of dynamic genes identified in shCon-transduced cells from clusters T5 and T9, was analyzed using GSEA and the scRNA-seq data (log_2_ fold change, Wilcoxon rank) to compare shCon versus shMll1-transduced cells. (**G**) Purified naive CD8 T cells were stimulated with plate-bound anti-CD3 and anti-CD28 (TCR) and chromatin-associated RNA (nascent transcription) was analyzed before and after stimulation. The heatmap depicts the log_2_ fold-changes in expression for the top 10^th^ percentile of dynamic genes that explained the transition potential of shCon cells in cluster T5 (i.e., not dynamic in shMlll cells), and that were also differential 48 hours after TCR stimulation (padj<=0.05), and were clustered by Euclidean distance. The expression of these genes in shMll1 versus shCon-transduced cells in the scRNA-seq data is also shown (*bottom* row). Chi-squared test for each row tests the fraction of genes that were upregulated against the fraction of genes upregulated in all genes in the given dataset. (**H**) Differential gene expression between shMll1 and shCon transduced cells in cluster T11 (log_2_ fold change, Wilcoxon rank) is displayed as a volcano plot. Genes highlighted in blue are significantly regulated genes within the TCF^hi^ gene set; those in red within the TE set.

### Mll1 promotes transcription of dynamic genes that drive development of nascent T_SCM_ precursors

*Tcf7*^hi^ T_EFF_ cells might comprise T_SCM_ cells that bypass full T_EFF_ cell differentiation during acute infection (*12, 16*), but the transcriptional control that specifies their early formation is still unclear. We anticipated that the effect of Mll1-deficiency on early developmental trajectories might clarify this process, and therefore used single-cell RNA-sequencing (scRNA-seq) to analyze shCon-transduced and shMll1-transduced P14 cells 5 days after LCMV_Arm_ infection in the context of cells isolated on days 3 and 8 pi that derived from separate transfers of small numbers of wildtype naive P14 cells. Cells from all time points separated into 13 Leiden clusters (Fig 3B, *top*), and were numbered according to increasing latent time to define a potential developmental order (fig S8A). In the unsupervised arrangement, transduced cells from day 5 pi were concentrated in clusters located between clusters that mainly comprised cells from days 3 and 8 pi, consistent with their development in actual time (Fig 3B, *bottom*). The enrichment of known gene expression signatures aided defining the biological nature of cells in each cluster (Fig 3C). Cells transduced with shCon or shMll1 contributed to the clusters differentially (Fig 3D), and exhibited distinct RNA velocity-derived transition probabilities (Fig 3E-F and fig S8B), indicating they propagated through the trajectory differently.

Cluster T1 comprised TCR-activated intermediates enriched with the signature of T_CM_ cells (Fig 3C) and exhibited transition potential into clusters T2 and T3 (fig. S8B). Cluster T2 comprised a small number of cells from day 3 pi that strongly repressed the *Tcf7*^hi^-effector cell signature (Fig 3C). In contrast, cluster T3 comprised a larger number of shCon-transduced cells from day 5 pi that upregulated the signatures of *Tcf7*^hi^-effector and T_prog_ cells (Fig 3C). Cells from cluster T3 branched into cluster T5 (fig S8B), an activated and unbiased T_EFF_-like state (Fig 3C), and into cluster T6 (fig S8B), a state positively enriched with signatures of both KLRG1^hi^ CD127^hi^ double positive (DP) effector cells, and tissue-resident memory (T_RM_) cells (Fig 3C). This suggests most activated cells initially express Tcf1-driven gene expression before effector cell specification occurs, consistent with prior studies (*49*). However, shMll1-transduced cells were virtually absent from cluster T3, and significantly fewer contributed to clusters T5 and T6 compared to shCon-transduced cells (Fig 3D). Residual shMll1-transduced cells from cluster T3 were negatively enriched Tcf1 and Tox-promoted gene expression and the signatures of *Tcf7*^hi^-effector and T_prog_ cells compared to shCon-transduced cells (fig S8C), and lacked transition potential into either clusters T5 or T6 (fig S8B). Thus, Mll1 was required for the development or stability of activated cell states that sustain T_SCM_-related gene expression.

Mll1 governed the transition potential of cells in cluster T5, which normally diverged into cell states that either sustained and upregulated the signature of *Tcf7*^hi^ effector cells, or that downregulated this signature and acquired effector-like signatures. Control-transduced P14 cells in cluster T5 had transition potential into clusters T7, T8, and T9, whereas shMll1-transduced T5 cells exhibited transition potentials into clusters T6 and T9 (Fig 3E and fig S8B). Thus, Mll1 normally promotes transition into clusters T7 and T8. Con-transduced cells in clusters T7 and T8 maintained the expression of both *Tcf7*^hi^ effector and T_prog_ cell signatures (Fig 3C, *Day 8 LCMV_Arm_*), upregulated genes expressed by memory and naive cells (Fig 3C, *naive & memory*, *IV*) and repressed gene expression of acutely activated cells (Fig 3C, *early activation genes, III*), whereas those in cluster T9 less strongly expressed the *Tcf7*^hi^ effector and T_prog_ signatures compared to clusters T7 and T8, maintained expression of early activation genes (Fig 3C, *early activation genes, III*), and upregulated the signatures of both DP effector cells and T_RM_ cells (Fig 3C, *Day 8 and Day 30 LCMV_Arm_* signatures). Conversely, shMll1-transduced cells in all three of these clusters less highly expressed genes normally induced by Tcf1 and Tox and overexpressed gene expression promoted by the TFs T-bet (*Tbx21*) and Blimp1 (*Prdm1*) (fig S8C-D). Thus, Mll1-deficient T5 cells did not normally establish T_SCM_-related gene expression, were effector-cell biased and preferentially diverged into effector-like states, suggesting this was a proximal defect of Mll1 deficiency.

To define how gene expression regulated the developmental potential of cells in cluster T5, the genes whose transcriptional behavior was most dynamic between clusters (driver genes) were extracted from shCon-transduced and shMll1-transduced cells. Driver genes explain the derived transition potentials of cells (*56*) in the trajectory and were substantially distinct in shCon-transduced and shMll1-transduced cells (table S4). The drivers in shCon-transduced cells identified from the entire trajectory (Fig S8E), and those specifically extracted from clusters T5 and T9 (Fig. 3F) were significantly enriched among genes that were more highly expressed in shCon-transduced cells, suggesting that their normal dynamics in the trajectory involved positive regulation by Mll1. In addition, driver genes specific to shCon-transduced cells from clusters T5 and T9, or common to both shCon-transduced and shMll1-transduced cells from both clusters (Fig S8F), were more highly expressed in shCon-transduced cells of cluster T5 compared to T9 (fig S8G, p=0.0002), whereas those exclusive to shMlll-transduced cells were more highly expressed in shCon-transduced cells from cluster T9 than in cluster T5 (fig S8G, p=0.0077). Together, these results suggest that Mll1 is required to activate driver genes whose dynamics explain early developmental transitions and to prevent activation of alternative driver genes that explain transition into later states. TCR stimulation was likely to positively regulate the dynamics of driver genes in cluster T5, because genes that are transiently induced 24 and 48 hours after acute infection were more highly expressed in cluster T5 cells (Fig 3C, *early activation II*, and *III*). In addition, nascent transcription of driver genes in shCon-transduced cells from cluster T5 were strongly regulated in naive CD8 T cells upon TCR and co-stimulation (Fig 3G). Thus, the transcriptional dynamics of genes which account for early transition potentials appear to be governed by TCR-signals and positively regulated by Mll1.

Cluster T11 appeared to be a distinct T_SCM_ precursor state because it was most strongly enriched with the signatures of *Tcf7*^hi^ effector and T_prog_ cells compared to all clusters in the trajectory, and had upregulated both naive and memory cell-associated gene expression including the T_CM_ cell specific signature (Fig 3C). Cluster T11 cells mainly derived from transduced cells on day 5 pi and bridged development into cell states from day 8 pi that first repressed the Tcf7^hi^-effector and T_prog_ signatures (cluster T12) and then transitioned into cluster T13 which activated TE cell gene expression (Fig 3E and fig S8B). Thus, T_EFF_ cell populations found in the spleen on day 8 pi developed directly from a distinct T_SCM_-like precursors on day 5 pi. Compared to shCon-transduced cells in cluster T11, gene expression in shMll1-transduced cells was negatively enriched with signatures of *Tcf7*^hi^ effector, T_prog_ and mature T_CM_ cells, but was positively enriched with signatures of TE and T_EM_ cells (fig S8C). Differential expression demonstrated that shMll1-transduced T11 cells expressed significantly less of the *Tcf7*^hi^ effector cell signature genes *Tcf7*, *Tox*, *Aff3*, *Bcl2, Ltb*, *Slamf6* and *Ctla2a* and significantly more of the TE cell signature genes *Klrg1*, *Gzma*, *Ccl5*, and *Cd48* (Fig 3H, p=0.0002 and p=0.0046 respectively). In contrast, there was no bias in the number of *Tcf7*^lo^ effector cell signature genes that were differentially expressed in shCon-transduced cells compared to shMll1-transduced cells, indicating that the reciprocal effects of Mll1-deficiency on *Tcf7*^hi^ effector and TE cell gene expression were specific (p=0.1522). These results indicate Mll1 stabilizes an early T_SCM_ precursor state and prevents premature activation of TE cell gene expression, which explains the altered frequency distributions of TE and MP cells that develop from shMll1-transduced cells on day 8 pi (Fig 2D).

### Mll1 pioneers widespread and T_MEM_-specific H3K4me3 deposition at TSSs during TCR stimulation

The mechanics that program enhanced transcriptional competence in T_MEM_ cells compared to naive cells are still unknown. Histone H3 lysine 4 tri-methylation (H3K4me3) deposition near transcription start sites (TSSs) defines poised and active promoters (*57*), and may positively regulate transcriptional elongation (*58*). Many genes lacking H3K4me3 in naive CD8 T cells acquire this modification in mature T_EFF_ and T_MEM_ cells (*31, 59*), suggesting it is a key chromatin regulatory step that differentiates T_MEM_ cells from their naive ancestors. Although Mll1 is the archetype H3K4-trimethylase (*60, 61*), its disruption in mammals has only demonstrated reductions in H3K4me3 at limited number of genes and other COMPASS-family methyltransferases are thought to establish and maintain the majority of H3K4me3 (*43, 62, 63*). However, we found that during TCR stimulation the relative abundance of H3K4me3, and RNA Polymerase II (Pol II) hyperphosphorylated on serine-5 (Ser5-P) or serine-2 (Ser2-P) C-terminal domain (CTD) residues (fig S9A) increased in conjunction with expression of the catalytic component of MLL-COMPASS-like complexes (Mll1-C), suggesting that Mll1 could promote transcriptional reprogramming during initial naive cell activation.

ChIP-seq analysis demonstrated that regions of H3K4me3 deposition (“peaks”) in naive and TCR-stimulated cells overlapped extensively with those present in *ex vivo* T_MEM_ (KLRG1^lo^ CD127^hi^) cells (*31*), indicating that naive and antigen-experienced CD8 T cells share an extensive core H3K4me3 landscape (*31, 59*) (fig S9B). In addition, TCR stimulation remodeled the naive cell landscape by transiently reducing H3K4me3 at some peaks relative to naive cells and progressively increasing *de novo* H3K4me3 peaks, which increasingly overlapped those present specifically in *ex vivo* T_MEM_ cells (Fig 4A, fig S9B). Notably, approximately 20% of all peaks that exhibited significantly greater H3K4me3 in T_MEM_ cells relative to naive cells were induced within the first 72 hours of TCR-stimulation (fig S9B). Thus, a substantial fraction of the T_MEM_ cell-specific H3K4me3 landscape is established during initial TCR stimulation, and is potentially maintained “epigenetically” as some activated cells develop into T_MEM_ cells.

**Fig 4:**
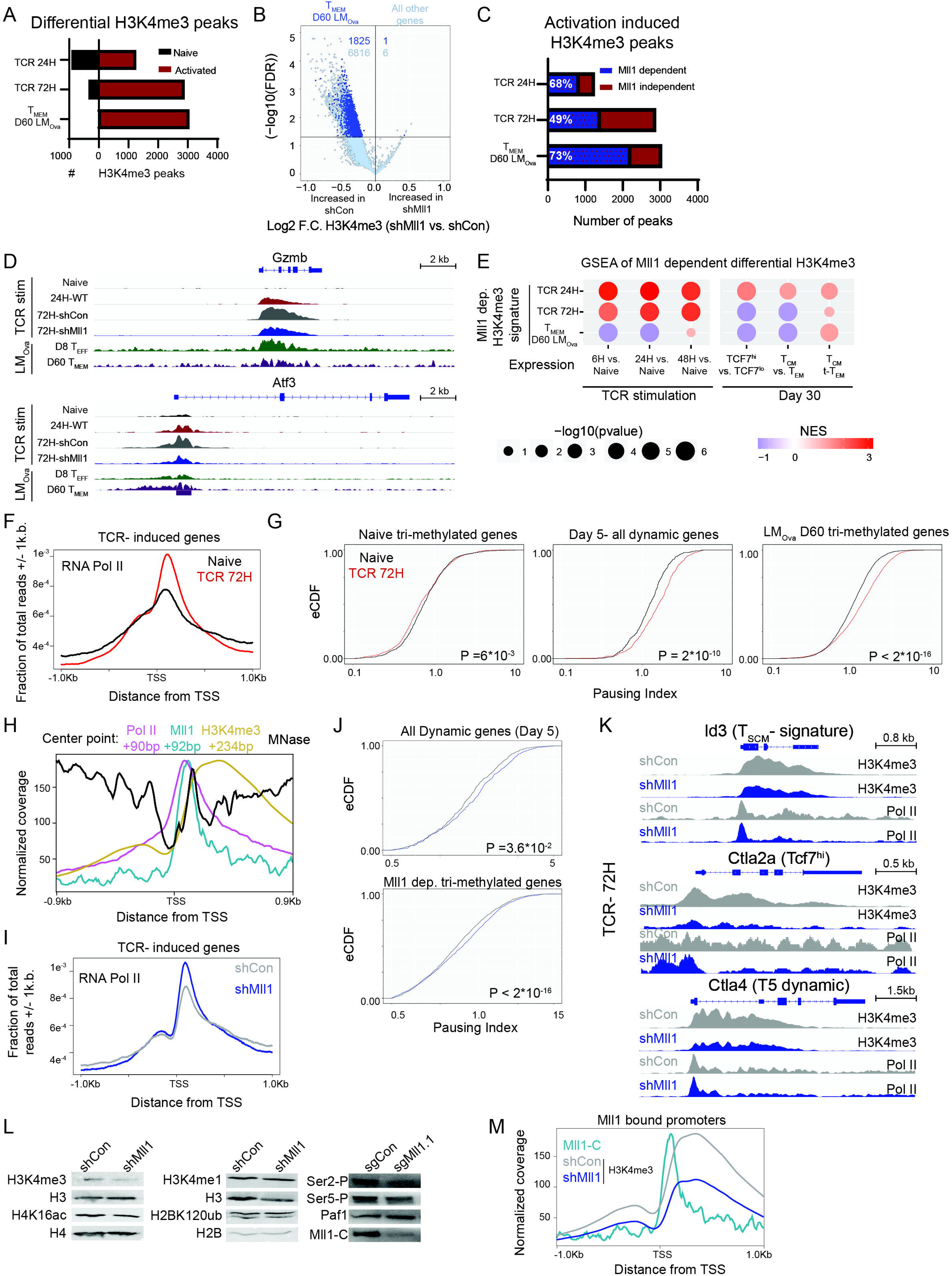
TCR signals and Mll1 establish T_MEM_ cell-specific H3K4me3 deposition and RNA Pol II pausing. **(A)** Bar chart shows the number of differential H3K4me3 peaks in activated versus naive cells (log_2_ fold change, P <= 0.05). (**B**) H3K4me3 peaks were annotated to the nearest gene and the average log_2_ fold change and false discovery rate (FDR) was calculated per gene in shCon-transduced (left) versus shMll1-transduced cells (right) 72 hours after TCR stimulation. The number of genes (grey) and those harboring T_MEM_ specific H3K4me3 peaks (dark blue dots) that were differential (FDR < 0.05) are indicated. (**C**) Stacked bar chart shows the number and fractions (white numbers) of H3K4me3 peaks that form after naive cells have been activated (red) whose H3K4me3 signal is significantly greater in shCon compared to shMll1 transduced cells 72 hours after TCR stimulation (blue bar). (**D**) Genome tracks show TCR-dependent and Mll1-dependent H3K4me3 deposition in representative genes. Coverage normalized to sequencing depth, and track heights normalized across TCR stimulated, naive and LM_Ova_ samples independently. (**E**) Bubble plot quantifies enrichment of differential gene expression (Wald statistic) from the indicated comparisons within genes harboring annotated H3K4me3 peaks that exhibited significantly increased H3K4me3 density in TCR-activated or T_MEM_ (LM_Ova_ D60) cells compared to naïve cells that were also dependent on Mll1 for H3K4me3 depostion(signatures). The bound set contains all genes bound by Mll1. (**F**) The fraction of total Pol II reads from naive cells (black) and cells activated by TCR and co-stimulation for 72 hours (red) located +/-1 kb of the TSS was calculated per base pair and plotted around the TSSs of genes induced transcriptionally by TCR signals (48 hours TCR versus naive, chromatin associated RNA-seq, P adj. < 0.05). (**G**) The ratio of normalized Pol II counts in promoters (-50bp to +300bp) to counts in gene bodies (+301 to +600 bp) (pausing index) was calculated using Pol II data from naive (black) and TCR-stimulated (72 hours, red) cells. Pausing indices were plotted as empirical cumulative distribution functions (eCDFs, P values computed with Kolmogorov Smirnov test) for genes with greater H3K4me3 in naive cells (naive H3K4me3), all dynamic genes from day 5 pi (see Fig 3G, Day 5 dynamic genes), and genes with greater H3K4me3 in T_MEM_ cells compared to naive cells (LM_Ova_ D60 H3K4me3). (**H**) ChIP-seq and MNase-seq signals centered on Mll1-bound TSSs were plotted. Coverage was scaled across samples and calculated on a per bp level. (**I**) The fraction of total Pol II reads in shCon (grey) and shMll1 (blue) cells located +/-1 kb of the TSS of genes dependent on Mll1 for H3K4me3 deposition (FDR <= 0.05) were calculated per base pair and plotted. (**J**) The pausing indices of the indicated gene sets from shCon (grey) and shMll1 (blue) are plotted as eCDFs as in (G). (**K**) The deposition of H3K4me3 and Pol II is shown for representative loci in shCon (grey) and shMll1 (blue) tracks 72 hours after primary TCR stimulation. Coverage was normalized to sequencing depth. (**L**) Naive P14, or P14 Cas9-transgenic, CD8 T cells were transduced 16 hours after initial TCR stimulation with either shRNAmir-expressing retroviruses (*top*), or sgRNA-expressing retroviruses (*bottom*), respectively. Transduced cells were purified by FACS 72 hours after stimulation, equivalent cell numbers were solubilized in RIPA buffer and the resulting pellet (*left*) and supernatant (*right*) fractions were recovered and resolved by PAGE and analyzed by immunoblotting. (**M**) Binding of Mll1 (teal) and H3K4me3 from shCon (grey) and shMll1 (blue) at Mll1 bound TSS’s. Coverage was scaled to read-depth and calculated on a per bp level.

Genes that acquired H3K4me3 during TCR stimulation were more highly expressed in TCR-activated cells, and in both Tcf7^hi^ and Tcf7^lo^ T_EFF_ cells relative to naive cells, as well as in mature Tcf7^hi^ memory cells and T_CM_ cells, relative to t-T_EM_ cells (fig S9C-D). Thus, initial TCR stimulation induced H3K4me3 deposition at genes that become expressed in both T_EFF_ and long-lived T_MEM_ cells. Acute depletion of Mll1 during TCR stimulation significantly reduced H3K4me3 deposition at more than half (∼8,000 peaks) of all peaks identified in TCR activated cells (∼15,000 total peaks) (Fig 4B). This included ∼70%, and 49% of all *de novo* H3K4me3 peaks induced 24, or 72 hours after TCR stimulation, respectively, and ∼70% of all H3K4me3 peaks that are normally upregulated in mature T_MEM_ cells that comprised a mixture of T_CM_ and T_EM_ cells (KLRG1^lo^ CD127^hi^ D60, LM_Ova_ (*31*)) compared to naive cells (Fig 4C-D). Within activation-dependent H3K4me3 peaks, regions dependent on Mll1 for H3K4me3 during TCR stimulation were positively enriched among genes that were more highly expressed in TCR stimulated cells (Fig 4E, *left*). In addition, Mll1-dependent H3K4me3 regions induced 24 hours after TCR stimulation were specifically enriched in genes that were more highly expressed in T_CM_ cells (CD62L^hi^ CD127^hi^) relative to either T_EM_ (CD62L^lo^ CD127^hi^) or t-T_EM_ cells (CD62L^lo^ CD127^lo^) analyzed 30 days after LCMV_Arm_ infection (Fig 4E, *right*)(*64*); and those from T_MEM_ cells were positively enriched in genes more highly expressed in T_CM_ cells relative to t-T_EM_ cells (Fig 4E, *bottom right*). Thus, Mll1 was specifically required during initial TCR stimulation for H3K4me3 deposition in regions that define a large fraction of the T_MEM_ cell chromatin landscape, especially those specific to T_CM_ cells.

### Mll1 facilitates RNA Polymerase II elongation of dynamic genes during TCR stimulation

As expected, transcription positively correlated with H3K4me3 deposition in both naive and TCR-stimulated cells (Fig S9E). TCR stimulation increased H3K4me3 deposition and transcription of genes that were upregulated during the first 48 hours of infection, which included signature genes expressed in both Tcf7^hi^ and Tcf7^lo^ effector cells (Fig S9E, *top*). However, many genes that acquired H3K4me3 upon TCR stimulation did not immediately induce transcription, suggesting their full transactivation might depend on additional signals (Fig S9E). These predominantly comprised genes expressed in naive cells (blue) or genes that are upregulated in T_EFF_ or T_MEM_ cells at late times after infection (Fig S9E, *bottom*). Thus, TCR-induced H3K4me3 deposition correlated with immediate transcriptional activation, and established the potential for future transcriptional activation of T_MEM_-associated genes.

Analysis of total Pol II occupancy by ChIP-seq indicated that primary TCR stimulation induced widespread transcriptional pausing at genes that underwent H3K4me3 installation. Genes induced transcriptionally by TCR stimulation exhibited increased promoter and gene body-associated Pol II occupancy, compared to naive cells (Fig 4F and fig S10A). However, Pol II density at promoters increased more than in gene bodies, suggesting elongation of recruited Pol II became limiting upon TCR stimulation (fig S10A). Accordingly, Pol II pausing indices increased in genes that acquired H3K4me3 upon TCR stimulation (Fig 4F, fig S10B, table S5), in those whose transcriptional dynamics explained transition probabilities of cells in the trajectory analysis (Fig 4G, *middle*, table S5), and in those with greater H3K4me3 specifically in T_MEM_ (LM_Ova_ D60) cells compared to naive cells (Fig 4G, *right*, fig S10A, table S5). In contrast, genes marked with H3K4me3 specifically in naive cells exhibited a modest increase in their pausing indices in naive cells compared to TCR stimulated cells (Fig 4G, *left,* table S5). Thus, primary TCR stimulation induces H3K4me3 installation and Pol II pausing specifically in genes that establish the formation of T_EFF_ (LM_Ova_ D8) and T_MEM_ cells and that were Mll1-controlled. These results suggest that increased transcription of these genes is regulated by promoting release of paused Pol II (elongation).

To determine how Mll1 governed transcription, its binding was mapped in activated CD8 T cells using ChIP-exo5.0 (*65*). Mll1 primarily bound to the TSSs of active genes marked with H3K4me3, rather than promoter-distal hyper-accessible ATAC-seq regions marked with H3K4me1 (fig S10C). The summit of Mll1 occupancy localized 92bp downstream of TSSs (Fig 4H, *teal*) coincident with the summit of paused Pol II (Fig 4H, +90bp, *rose*), which was immediately upstream of an MNase-protected region corresponding to +1 nucleosomes (Fig 4H, *black*) and the summit of H3K4me3 density (Fig 4H, +230bp, *gold*). Mll1 signal was not enriched upstream of the TSS (Fig 4H, fig S10C). Thus, Mll1 was most likely recruited to paused Pol II near +1 nucleosomes after TCR activation. Depletion of Mll1 during TCR stimulation further increased promoter Pol II occupancy in the pause region (Fig 4I), resulting in increased Pol II pausing indices of dynamic genes (Fig 4J, *top,* table S5), and genes that required Mll1 for TCR-stimulation dependent H3K4me3 deposition (Fig 4J, *bottom* and 4K, table S5). Consistent with these results, depletion of Mll1 in TCR stimulated cells globally decreased both initiating (Ser5-P) and elongating (Ser2-P) forms of Pol II (Fig 4L)(*66*), suggesting Mll1 promotes transcription initiation and elongation. In line with these observations, shMll1-transduced cells exhibited a global decrease in molecule counts per cell in the scRNA-seq data (fig S10D), suggesting they manifested a broad defect in transcriptional activation in TCR stimulated cells *in vivo*. Thus, Mll1 most likely promotes efficient Pol II elongation upon TCR stimulation. Notably, Mll1-dependent deposition of H3K4me3 occurred immediately downstream of the Mll1 binding location (Fig 4M), suggesting that it might occur after Mll1-facilitated Pol II elongation events. Thus, Mll1 appears to have an early role in establishing H3K4me3 in T_MEM_-associated genes during the initial activation of naive CD8 T cells.

## Conclusion

Chromatin accessibility that develops in human CD8 T_EFF_ cells during the first week of live viral vaccination persists in slowly dividing T_MEM_ cells for nearly a decade (*11*), but the CRFs and transcriptional mechanics that establish and maintain these states are ill-defined. Our results demonstrate that Mll1 is essential for widespread H3K4me3 deposition during primary TCR stimulation, activation of T_SCM_ cell-associated gene expression and formation or stability of cells with T_SCM_-like phenotypes and functions during infections and tumors *in vivo*. In addition, we show that TCR stimulation and Mll1 regulate Pol II pausing in genes whose transcriptional dynamics *in vivo* drive early heterogeneity of activated CD8 T cell states during viral infection. These results are consistent with a model in which Mll1 configures Pol II at promoters for responsiveness to signals that govern Pol II pause release and transcriptional elongation to both specify early T_EFF_ cell developmental trajectories and to enable long-term capacitance of transcriptional activity in T_MEM_ cells. Notably, deposition of H3K4me3 at the 5’ ends of genes is known to occur as Set1-methyltransferases traffic with elongating polymerases (*67*). Our analyses indicate that Mll1 could traffic with Pol II to directly deposit H3K4me3, or potentiate the action of additional methyltransferases by facilitating modification and release of paused Pol II. In either model, Mll1 pioneers early H3K4me3 deposition during TCR stimulation that initially establishes the chromatin architecture of transcriptionally poised and active genes that define long-lived T_MEM_ cells.

## Acknowledgments

We would like to thank the Karbstein, Solt, and Sundrud labs for reagents and productive conversations, the Nettles lab for helping develop the E0771 model, and Dapeng Wang for performing initial MNase experiments. We would also like to thank the Flow Cytometry, the Genetic Perturbation Screening, and Genomics Cores at the Wertheim UF Scripps Institute for resources and services.

## Author contributions

Conceptualization: AJG, MAF, JJM, TV, HD, and MEP

Data curation: AJG, MAF, JJM, TV, and HD

Formal analysis: AJG, MAF, JJM, TV, HD, SDN

Funding acquisition: AJG, SC, AWG, and MEP

Investigation: AJG, MAF, JJM, HD, CT, SDN, DSA, SMT, and JK

Methodology: AJG, MAF, JJM, SB, SMT, and MEP

Project administration: AJG and MEP

Resources: SB, SC, AWG, and MEP

Software: AJG, TV, HD, and SDN

Supervision: SC, AWG, and MEP

Validation: AJG, MAF, JJM, and TV

Visualization: AJG, MAF, JJM, TV, and MEP

Writing original draft: AJG and MEP

Writing review and editing: All authors

### Vectors, shRNAmirs, CDNA constructs and retroviral packaging

The MSCV-based retroviral vectors MigR1, LMPd, and LsgA were used to transduce activated CD8^+^ T cells. MigR1 encoding IRES-GFP, was used to deliver *RUNX3* (long form) CDNAs(*55*). LMPd, encoding Pgk-Ametrine1.1 or Venus was used to express shRNAmirs, whose sequence was predicted by the shERWOOD algorithm and were purchased from TRANSOMICS(*69–71*). The LsgA vector was previously described and the Chop-Chop program was used to design guides(*72*). Retroviral supernatants were generated in Platinum-E packaging cells as previously described(*69*).

### Mice

C57BL/6J mice used in this paper were obtained from the Jackson Laboratory. P14 Thy1.1^+^ mice were a gift from R. Ahmed (Emory University). Thy1.1^+^,Thy1.2^+^ positive mice were generated by crossing the P14 mice to wildtype BL/6J mice. The Tcf7 reporter mice were purchased from Jackson and crossed to P14 mice to create P14-Tcf7 mice (*73*).The P14 TCRa^-/-^ were initially purchased from Taconic(*38, 74, 75*). All mice were maintained in specific-pathogen free facilities and used according to protocols approved by the Institutional Animal Care and Use Committee of TSRI-FL.

### Naïve T-cell culture and transduction

CD8^+^ T cells were isolated from naïve mice using negative selection magnetic beads (StemCell). For primary TCR-mediated activation, purified CD8^+^ T cells were plated at a density of 4x10^5^ cells/cm^2^ at a concentration of 1x10^6^/mL and on plates pre-coated with goat anti-hamster capture antibody and media supplemented with hamster anti-CD3 (2C11) and anti-CD28 (37.51) (1ug/mL each). For retroviral transductions, 16 hours after initial plating, the T cell conditioned media was taken off, saved, and replaced with retroviral supernatants containing 10ug/mL polybrene, and the cultures were immediately centrifuged at ∼ 500xg for 60 min at ∼ 37 degrees C, followed by incubation for an additional 3 hours at 37 degrees C at 10% CO_2_ incubator. Media containing retrovirus was aspirated and replaced with the saved conditions media. Cells were either transferred into mice (see infections) or continued to be cultured. For *in vitro* culturing, after 48 hours of total TCR stimulation, the activated cells were directly resuspended, counted, and re-cultured by diluting the cells into fresh media containing 10 or 100 U/mL rhIL-2 (NCI). Cells were re-cultured as 5x10^5^ cells/mL, every 24 hours following removal from the TCR stimulus(*38, 69*).

For experiments where cells were CTV labeled, after isolation, cells were pulsed with CTV for 10 minutes prior to activation.

### Screening Analysis

The geometric means from the live singlets transduced with shRNAmirs were measured via FACs. Control RNAis were used as the population reference to calculate individual Z scores at day 6 while the full population was used at day 3. From these Z scores a per shRNAmir percentile shift was calculated and averaged across all shRNAmirs targeting the same gene. These average shifts were used to calculated Z-scores, pvalues, and RNAi effects, which was calculated as *z* * (1 - *pvalue*).

GSEA was computed on the RNAi effect values using cluster profiler(*76*) (Subramanian, 2005 #485}. Chromatin regulator family gene sets were custom built for this paper

### Pathogens, infections, and colony forming assays

LCMV-Armstrong and LCMV-Clone13 were produced as previously described (*77*). For experiments using in vitro transduced P14 cells, 50,000 activated cells were adoptively transferred into naïve C57/Bl6J mice r.o and an hour later infected with either 2×105 PFU of LCMV-Arm i.p. or 2x106 PFU r.o. For Listeria monocytogenes-GP33 (LM-GP33) was grown as previously described on brain heart infusion media(*78*). Mice were infected with 10,000 CFU for recall experiments.

### Tumor transplant, ACT, and harvesting

E0771 cells containing either a control MigR1-GFP or MigR1-GP33-GFP were cultured per providers instructions. For transplant, 1x10^6^ tumors cells per mouse were resuspended in 100μL of media and mixed 1:1 with 100uL of high concentration growth factor reduced (HC-GFR) matrigel (Corning) on ice and injected s.c. into mice mammary fat pads. 2 weeks later, 1x10^6^ P14 cells transduced with shRNAmirs were transferred into tumor bearing mice.

For harvesting, tumors were excised and dissociated into ∼1cm pieces. Tumor were digested for 10 minutes in 10uL/mL DNAseI and 1uL/mL Collagenase and strained over a 70μM filter and spun for 10 minutes at 300g. Supernatant was then aspirated, and pellet was resuspended in 1mL of ACK lysis buffer for 5 minutes with occasional agitation. After 5 minutes, lysis buffer was diluted with 5 mL of cTCM and strained over a 45 μM filter and spun for 10 minutes at 300g. Pellets were then resuspended and spun in a 60/40 Percol gradient to specifically enrich lymphocytes.

B16 ACT experiments were performed as previously described(*79, 80*).

### Ex-vivo isolation from tissues-spleen

For EdU experiments, mice were pulsed for either four hours (day 5) or overnight (day 8) with Edu before sacrifice.

T cells from mice infected with LCMV before day 6 were digested as previously reported(*81*). In brief, spleens were harvested into T-cell optimized media (cTCM)(*38*) and cut into ∼1cm pieces. Spleens were digested for 10 minutes in 10uL/mL DNAseI and 1uL/mL Collagenase. Spleens were then treated as all other splenic experiments.

Spleens were strained over a 70μM filter and spun for 10 minutes at 300g. Supernatant was then aspirated, and pellet was resuspended in 1mL of ACK lysis buffer for 5 minutes with occasional agitation. After 5 minutes, lysis buffer was diluted with 5 mL of cTCM and strained over a 45 μM filter and spun for 10 minutes at 300g. For analytic flow cytometry, pellet was resuspended in 1mL of FACS buffer (1xPBS, 2% FBS, 10mM EDTA).

For sorting, cells were negatively isolated using a modified StemCell CD8^+^ kit. In brief, biotinylated αCD19, αCD4,αB220,and αTER-20 antibodies were added to cell pellet in 1mL of cTCM and 50μL of rat serum. After 10 minutes, the manufactures instructions and reagents were followed and used. Cells were then transferred into FACS buffer for staining at ∼30-40 million cells per mL (See staining protocol). After staining, cells were sorted via a BD Aria Fusion sorter.

### Flow cytometry

After tissue isolation, cell numbers were counted via an Acuri C6 or MACSQuant 10. Counts were normalized across individual mice to the highest number of P14 cells to reduce experimental variability. Samples were transferred into FACS buffer and stained for either 20 minutes at RT or 40 minutes at 4C. Samples were then washed and fixed for 10 minutes at 4 C in 4% PFA. Samples were washed again and placed into FACS buffer at 4 C until acquisition. Samples were acquired on a BD LSR2 or a Cytek Aurora

### Generation of bulk RNA-seq libraries and sequencing

For bulk RNA-seq analysis of *in vivo* CTL, distinct phenotypic subsets were isolated by FACS and sorted into FBS coated DNA LoBind Eppendorf 1.5mL with 200uL of 1xPBS, 1mM DTT, and 5U of Recombinant RNAse inhibitors (Takara), and then were washed in 1x PBS. Full-length CDNAs were generated directly from ∼10,000 cells per sample using oligo(dT) priming and the SMART-Seq v4 Ultra Low Input RNA Kit (Clontech), and 150 pg of CDNA was used to prepare strand-specific paired-end sequencing libraries (Nextera XT, Illumina). Libraries were quantified using Qubit HS DNA assay (Invitrogen) and their quality was assessed on Agilent Technology 2100 Bioanalyzer using a High Sensitivity DNA chip (Agilent) and dual-indexed pooled libraries were sequenced. Reads were aligned with salmon using validateMappings(*82*). Aligned reads were passed to DESEQ2 for differential gene analysis(*83*). All gene sets used were derived from previously published papers: EMP, naïve, and 48H(*1*); TE,MP,EE, and DP(*55*) ; T_RM_, T_CM_, and T_EM_ (*68*) Tcf7(*12*); and T_prog_(*84*).

Nascent RNA-seq library preparation, sequencing, and data analysis were previously described(*1*)

### Single cell clustering, cluster specific gene expression, and pseudotime analysis

Single cell library preparation, sequencing, and alignment were previously described(*1*).

Single cell data was manipulated and analyzed first in the scanpy environment (*85*). Cells from D5 shCon, D5 shMll1, and D8 were down sampled to 800 cells and those matched to D3 to 400 cells. Mutual nearest neighbor batch correction was used to remove batch effect from the two separate sequencing experiments (mnn_correct) (*86*). Using the merged experiments, principal components (tl.pca) and nearest neighbors were calculated (pp.neighbors, neighbors=100, pcs=25) and then used to create Leiden clusters (tl.leiden, resolution =1). These clusters were then used to create a PAGA graph to estimate and simplify the connectivity of the clusters (tl.draw_graph) (*87*).

Data from these clusters was used for all single cell differential expression calculations using the Wilcoxon method (tl.rank_genes_groups). For GSEAs the score for each given analysis was used as the rank ordered input. To calculate pseudotime, an arbitrary D3 cell from Leiden cluster 7 (T1 in the text) was selected as the initial cell (tl.dpt).

### RNA velocity, latent time, transitions, and dynamic genes

To calculate RNA velocity, scvelo was used(*56*).First, using the pre-pseudotime PAGA analysis, neighbors, moments and initial velocity were calculated (pp.neighbors neighbors =100, pcs=50, pp.moments, and tl.velocity respectively). Next dynamic genes were recovered and used to compute the dynamical model (tl.recover_dynamics, tl.velocity). Using these dynamic genes both latent times (tl.latent_time) and ranked dynamic genes (tl.rank_dynamical_genes) were calculated. To calculate individual dynamic genes for either control or Mll1 deficent cells, cells originating from either Mll1 deficent or control samples were removed prior to scVelo analysis. These dynamic genes were then used to calculate transition probabilities.

**Chromatin I.P**: T cells were fixed in PBS containing a 1:10 ratio of fixation buffer (11.1% V/V MeOH free formaldehyde, 100mM NaCl, 1mM EDTA, and 50 mM Hepes pH7.5) for 15 minutes on ice. Fixation was quenched with a 1:20 addition of 2.5M glycine for an additional 15 minutes. Cells were then washed with PBS and nucli were extracted (10mM Tris pH7.5, 10mM EDTA, 150 mM NaCl, and 1% TritonX-100). Nucli were resuspended in lysis buffer (50mM Tris pH 8.0, 150mM NaCl, 1mM EDTA, 1% Nonidet P-40, 0.1% SDS, and 0.5% Deoxycholate, supplemented with protease and phosphate inhibitor tablets (Roche)). To make mono and di nucleosome size fragments for I.P, sonication was performed in covaris mini tubes for 30-45 minutes with following settings: peak power-160, duty factor -10, cycles/burst=200.

Before the addition of antibody, chromatin was brought up to 1mL total volume in lysis buffer and pre-cleared via the addition of 50uL of Protein G Dynabeads per 10^7^ nucli equivalents at 4C for 2-3H. 5% of each reaction was then saved for making the input control. The equivalents of 10-20 million nucli were subdivided for each I.P per sample and brought up to 1mL total volume. Antibodies were then added to the chromatin and I.P’d for ∼16-20H at 4C. Each sample was then conjugated to fresh Protein G Dynabeads (5uL beads per 1ug of antibody) at RT for 15 minutes. After conjugation cells were washed twice with lysis buffer and once with TE. Decrosslinking than occurred via 30 minute treatment with RNAseA (100ug/mL) at 37C, followed by overnight digestion of protein via proteinase K (1mg/mL) at 65C(*88*).

For high resolution ChIP-Exo experiments, samples were washed as above and then ChIP-Exo 5.0 was followed as previously reported(*88*).

### Library preparation and sequencing

DNA from chromatin preps was extracted via phenol: chloroform extraction and then purified and concentrated via EtOH cleanup. Resulting DNA was quantified via Qubit. 1 to 5 ng of DNA were used to create sequencing libraries via NEB Ultra II DNA Kits. After amplification, resulting libraries were checked for size (∼300bp) via either Bioanalyzer 2100 or TapeStation 4200. Samples were then pooled at equal molar amounts of library and concentrated/ size selected via AMPure beads. Library was then run on NEXT-Seq 500.

### ChIP-Seq read processing and peak calling

Reads from sequencing were trimmed using trim_galore (--paired --length 24 --stringency 3) and aligned to GRCm39 using bowtie2 (--no-mixed –no discordant)(*89*). Aligned reads were then trimmed for duplicates and quality using samtools (samtools view -f 2 -q 20 -h 473-1.sam | samtools sort -n -o $QualFilt -O bam; samtools fixmate -m $QualFilt $fixmate; samtools sort -O bam -o $fixsorted $fixmate; samtools markdup -r $fixsorted $dupremoved)(*90*). Peaks were then called both per replicate and for all replicates via macs 2(callpeak -t $duprembam -c $controlbam --broad -f BAMPE -g mm -q 0.05 -B)(*91*)

### ChIP-Seq data processing

To determine significant differences between given peaks, the r package DiffBind was used(*92*). In the context of replicate controlled samples (shCon/shMll1 H3K4me3 and RNApol2) an additional replicate factor was added as part of the design. Peaks with a pvalue <=0.05 were considered significant for this analysis.

To determine gene enrichment, peaks from the given sample were then annotated to the nearest gene using ChIPseeker (*93*). The number of these genes that overlapped with previously described gene sets were counted and the simple fraction of the whole of the gene set was visualized using ComplexHeatmap (*1, 2, 12, 45, 55, 80, 94-96*).

To compute per gene log2FCs of H3K4me3 binding, the reported peaks from Diffbind were collapsed to gene and log2fcs from multiple peaks per gene were averaged. (*92*). Using these and the Log2FCs computed via DESEQ2 for the previously published naRNA-seq, data values were plotted on a two way dot plot(*1, 83*). Genes were marked by their presence in previously described gene clusters that were differential across time(*45*).

To compute the enrichment of one set of peaks within another, bedtools was used to determine the number intersecting peaks. The fraction of intersecting peaks over total number was then computed.

To calculate the pausing rate we adapted a previously published strategy (*97*). Annotated genes were separated into their promoters (TSS +300,-50b.p) and proximal gene body (301-600 b.p) and the number of reads in either region were calculated. These were converted to transcripts per million and the sum of TPMs to promoter or gene body were calculated and averaged across replicates. The per gene promoter sum and the per gene body sum were then divided to form the pausing index. Differences between indices were calculated via KS test.

## Supplementary Figures

**Fig. S1.**
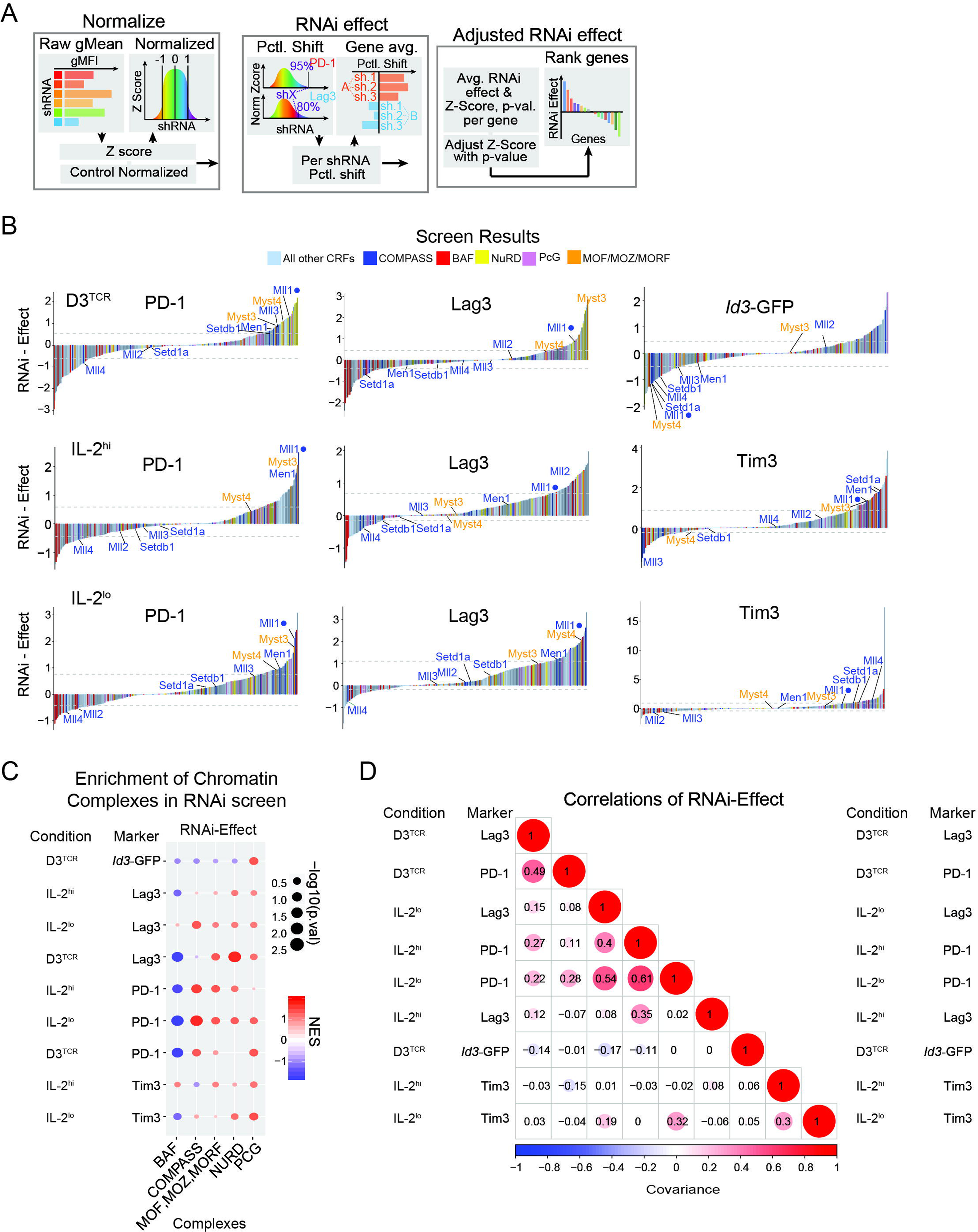
Chromatin regulators have both distinct and convergent effects on co-inhibitory gene expression. (**A**) Analytical method to compute and rank the average RNAi-induced effects (RNAi-effects) on marker protein expression in the screens that resulted from suppression of the same target gene with multiple distinct shRNAmirs. (**B**) Ranked RNAi-effects of each CRF in each screen are displayed as bar charts. Dashed lines indicate top and bottom 15^th^ percentile of RNAi-effects. (**C**) GSEA shows the enrichment of genes encoding subunits that assemble in biochemically defined chromatin regulatory complexes CRFs (bottom) within RNAi-effects on marker proteins in each screen (left). RNAi-effect scores were used to calculate enrichment. (**D**) Correlogram shows relationship of CRF RNAi-effects on different protein markers in distinct culture conditions by computing the correlation between RNAi-effect scores for each marker protein in different culture conditions. Size and color of circles in each box denote correlation intensity and directionality.

**Fig. S2.**
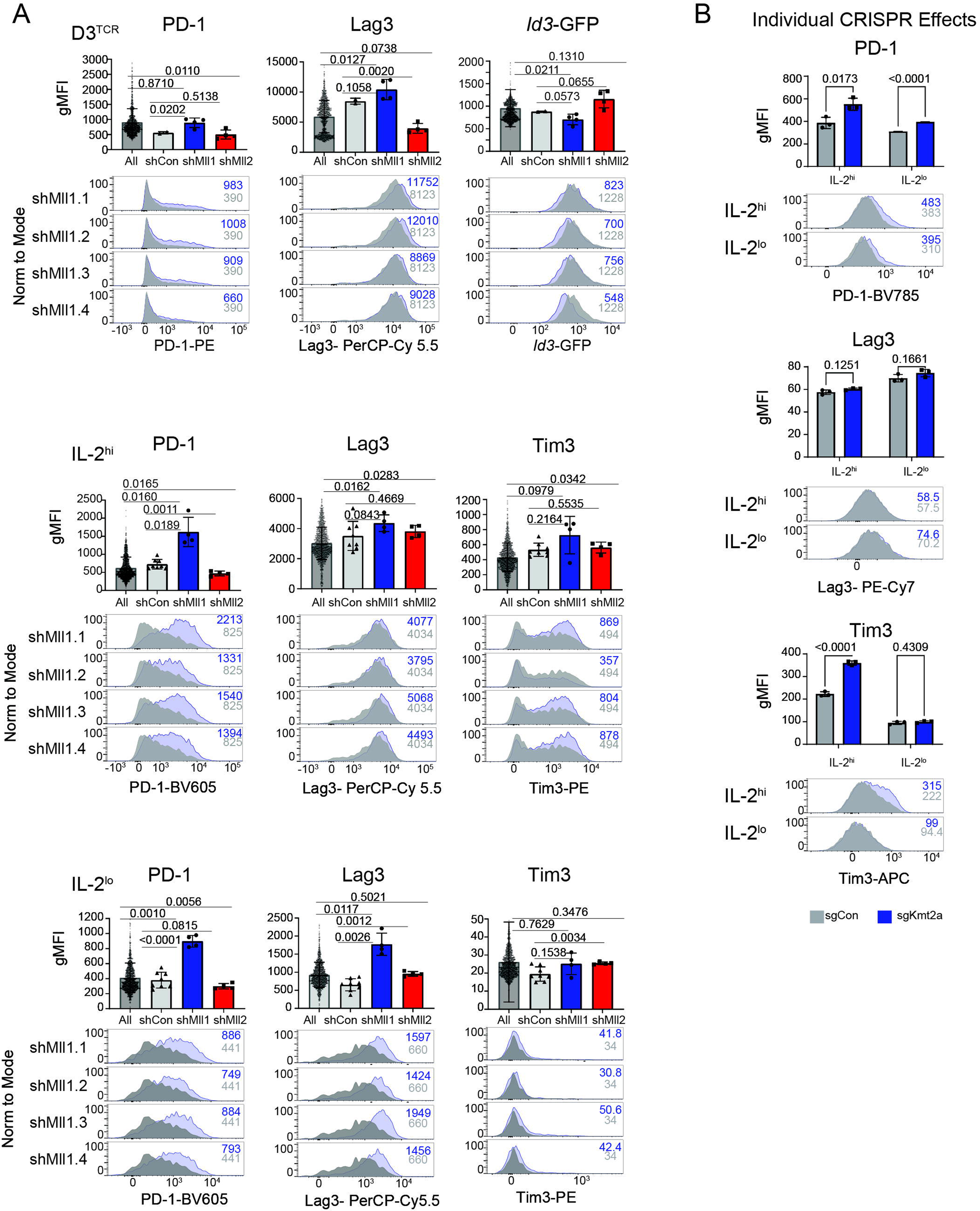
Mll1 deficiency coordinately decreases Tcf7 expression and increases co-inhibitory receptor expression. (**A**) Bar charts (*above*) indicate the mean expression of protein markers (gMFI) in cells transduced with all shRNAmirs specific for individual CRFs (or control genes) and for all shRNAmirs in the screen. Symbols indicate the individual raw gMFI values of cells transduced with unique shRNAmirs. P values were calculated using Welch’s t-test. Histograms (*below*) show representative flow cytometry data for individual shRNAmirs specific for control genes or Mll1 in each screening condition. Numbers in plots indicate gMFI values. (**B**) Bar charts (*top*) and representative histograms (*bottom*) show marker protein expression in P14 Cas9-transgenic CD8 T cells transduced with sgRNAs targeting *Kmt2a* (symbols, n=3). Numbers in plots indicate gMFI.

**Fig. S3.**
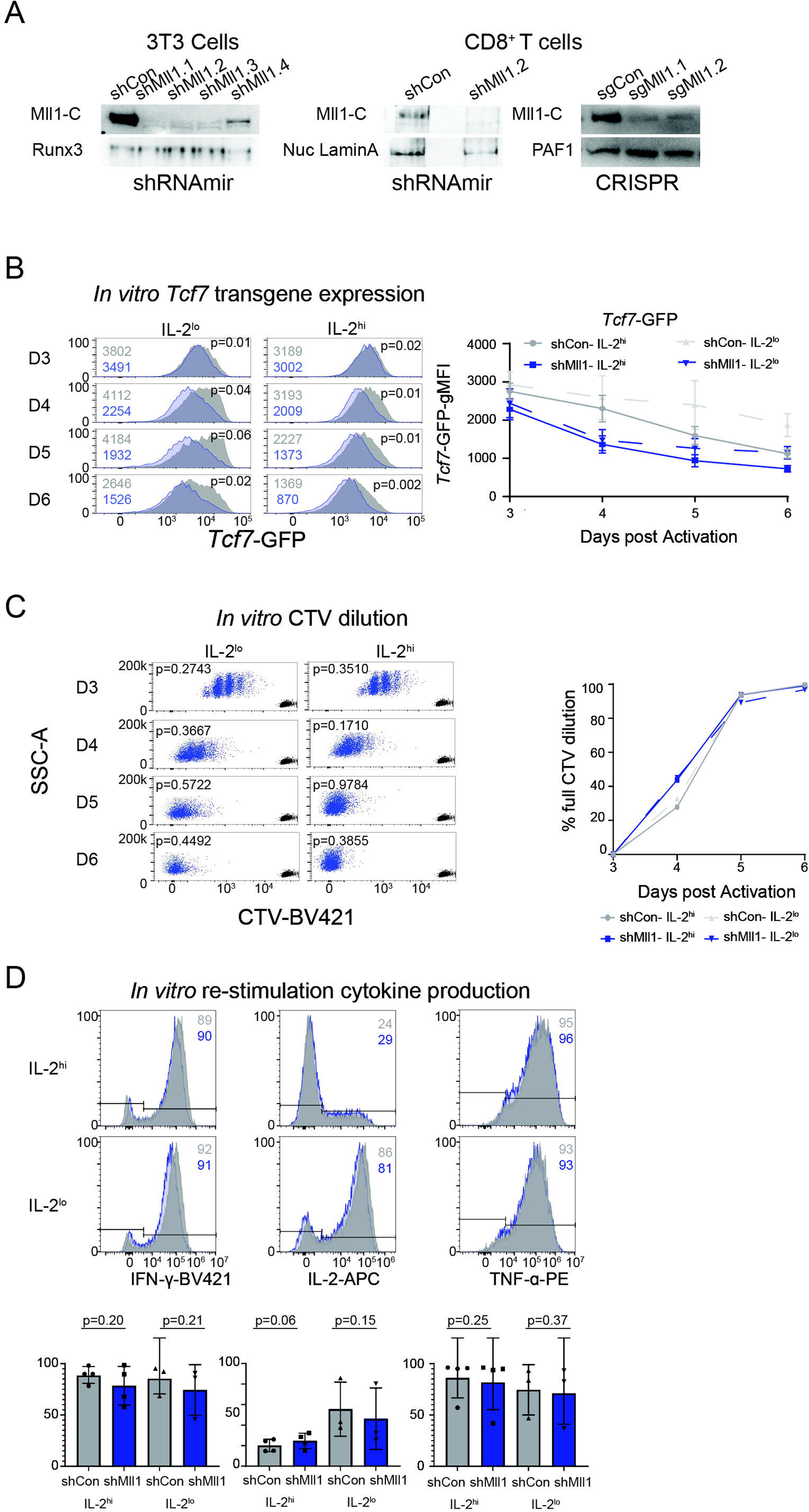
Mll1 deficient cells are not generally defective. (**A**) Mll1 expression was determined in whole cell lysates prepared from CD8 T cells transduced with either shRNAmirs (*left*) or sgRNAs (*right*). Transduced cells were purified by FACS three days after TCR activation and transduction. (**B**) Representative flow cytometry histograms from three biological replicates show *Tcf7*-GFP expression in P14*^Tcf7^*^-GFP^ cells each day in culture after transduction with shCon or shMll1 shRNAmirs (*top*). The gMFI (*left*) and P values (*right*, paired t-test, comparisons from three biological replicates) are indicated in each plot and are summarized in the line graph (*bottom,* n=3). (**C**) Histograms show CTV labeling of naive P14 CD8 T cells (black) and its dilution after gating on cells transduced with shCon or shMll1 shRNAmirs each day after activation (*left*). Numbers on the plot show P values (paired t-test, from three biological replicates) after comparing the percent of shCon to shMll1 cells that fully diluted CTV, which is summarized in the line graph (*right*). (**D**) Histograms show cytokine expression detected by intracellular staining in cells transduced with control or Mll1 specific shRNAmirs after 6 days of culture in IL-2^lo^ or IL-2^hi^ conditions, followed by 4 hours of re-stimulation with phorbol 12-myristate 13-acetate (PMA) and ionomycin. Numbers on the plot show the percent of cytokine positive cells (*top*) and the bar charts (*bottom*) summarize data as the means from four replicate experiments (P values, unpaired t-test).

**Fig. S4.**
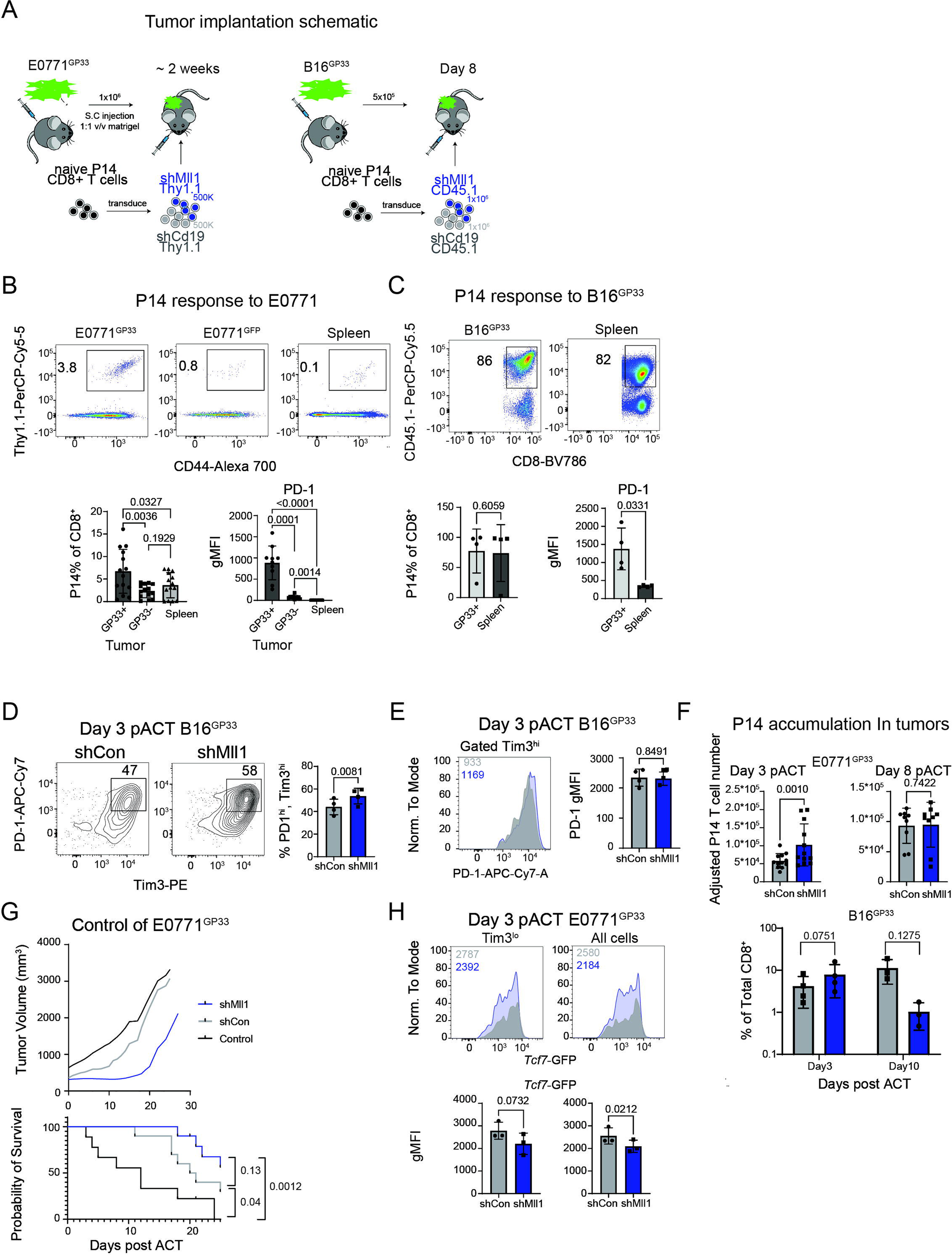
Mll1 restrains co-inhibitory protein expression and promotes formation of T_prog_ cells in the tumor microenvironment. (**A**) Schematics for the T cell adoptive transfer models using E0771 triple negative breast cancer (TNBC) (*top*) and B16 melanoma (*bottom*) tumors. (**B-C**) Representative flow cytometry plots from cells in mice transplanted with E0771^GP33^ and E0771^GFP^ in contralateral mammary fat pads after gating on CD8+ cells from different tissues (B, n=3), or B16^GP33^ flank tumors (C, N=4), as well as their matched spleens (*top*). Numbers show percent of donor Thy1.1+ CD44+ (P14) cells (B) or percent of CD45.1 among gated CD8+ cells (C). Bar charts (*bottom*) summarize the mean percentage of total CD8+ cells and gMFI of PD-1 expression on P14 cells from mice (symbols) bearing E0771 (B) or B16 (C) tumors, and were compared using paired t-tests. (**D**) Representative flow cytometry plots after gating on P14 CD8^+^ T cells transduced with shRNAmirs specific for negative control genes or Mll1 in B16^GP33^ tumors, and comprise concatenated data from multiple mice (n=3). (**E**) Representative surface expression of PD-1 among Tim3^hi^ P14 cells after gating on shCon or shMll1 transduced cells and is summarized by the bar charts as the means from individual mice (symbols) in multiple experiments. P values were calculated via paired t-test. (**F**) Bar charts show numbers of transduced P14 cells within E0771^GP33^ (top) or B16^GP33^ (bottom) tumors. The absolute numbers were normalized for E0771^GP33^ at day 3 (left) or day 8 (right) pACT (Wilcoxon ranked-test). Accumulation in B16^GP33^ bearing mice was calculated as the percent of CD8+ within B16^GP33^ tumors at day 3 (*left*) or day 8 (*right*) pACT (paired t-test). (**G**) Line graph (top) shows control of E0771^GP33^ tumors in recipients that received no adoptively transferred P14 cells (black), or P14 cells that were either shCon (grey) or shMll1 (blue) transduced. Each line shows the average growth of tumors smoothed over five days. Kaplan Myer curve (bottom) shows mice within this experiment reaching 1,500mm^3^ tumor size. Significance was calculated using Mantel-Cox test. (**H**) Representative Tcf7-GFP expression in P14*^Tcf7^*^-GFP^ CD8^+^ T cells isolated from E0771^GP33^ tumors after gating on cells transduced with shCon and shMll1 shRNAmirs (concatenated data from multiple mice, n=3). The gMFI values are indicated in each plot and their means for all mice (symbols) are summarized in the bar charts. P values calculated via paired t-test.

**Fig. S5.**
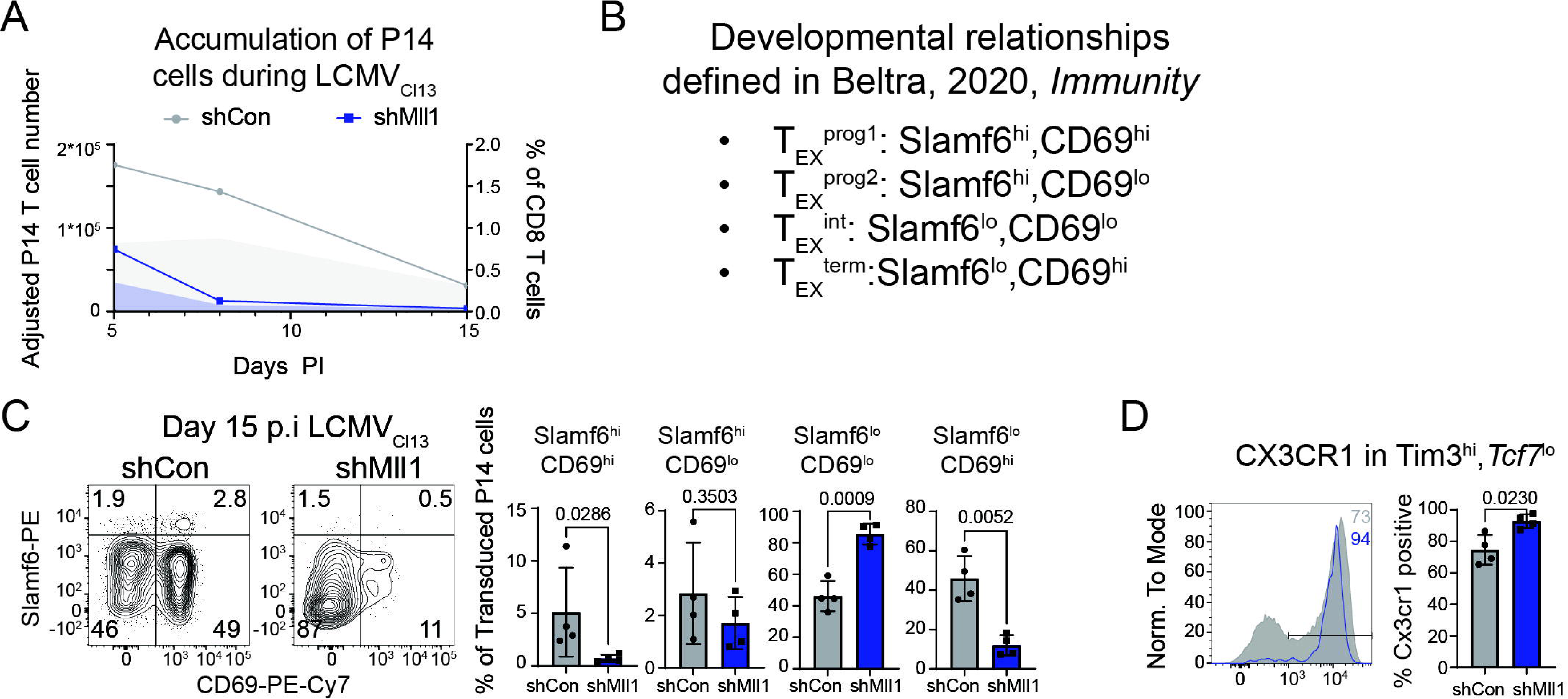
Mll1 is necessary for persistence and formation of T_prog_ cells during chronic infections. (**A**) P14 cell accumulation in response to LCMV_Cl13_. Symbols indicate normalized transduced P14 cell numbers in the spleen. Shaded portions of the plot indicate the normalized percent of transduced P14 cells of all CD8^+^ cells. Both data present the mean of n≥3 individual mice. (**B**) Summary of T_EX_ development markers from Beltra, 2017, *Immunity*(*26*) (**C**) Flow cytometry plots show representative staining after gating on P14 CD8^+^ T cells transduced with shCon and shMll shRNAmirs and comprise concatenated data from multiple mice (n=4). Bar charts summarize the mean frequencies of phenotypic T_EX_ cell populations from individual mice (symbols). P values calculated via Welch’s t-test. (**D)** Representative expression of CX3CR1 within gated Tim3^hi^Tcf7^lo^ P14 cells comprises concatenated data from multiple mice (n=4). Bar charts summarize the means of individual mice (symbols). P values calculated via Welch’s t-test.

**Fig. S6.**
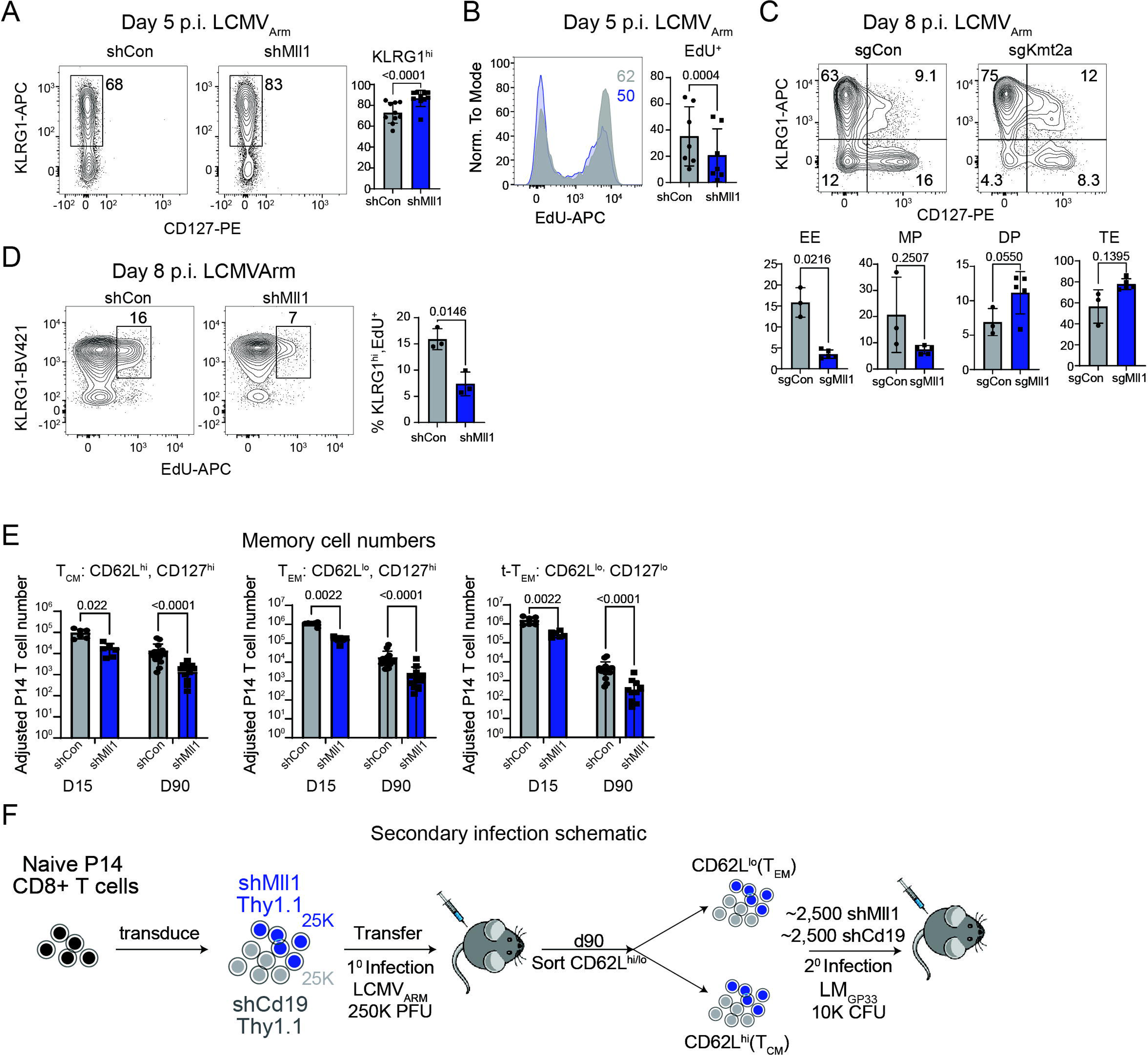
Mll1 promotes proliferation of responding T_EFF_ cells during primary infection. **(A-D)** Representative flow cytometry of transduced P14 CD8^+^ T cells in the spleen after LCMV_Arm_ infection are data concatenated from 2-5 mice. Values in the bar plots summarize the mean frequencies of the indicated phenotypic populations for all mice (symbols). P values were calculated with paired t-tests. (A) Representative KLRG1 staining on transduced P14 cells. (**B**) EdU incorporation is shown in the histogram after gating on P14 cells transduced with shCon or shMll1 shRNAmirs. (**C**) Representative staining of KLRG1 and CD127 on P14-Cas9 CD8 T cells transduced with sgCon or sgKmt2a specific sgRNAs were analyzed on day 8 pi in the spleen. (**D**) EdU incorporation was measured as function of KLRG1 expression on day 8 pi, after gatining on P14 cells transduced with shCon or shMll1 shRNAmirs. (**E**) The absolute numbers of T_CM_, T_EM_ and T_t-Tem_ cells in the spleen after gating on shCon and shMll transduced P14 cells were quantified for multiple mice (symbols) 15 and 90 days after LCMV_Arm_ infection. P values were calculated by Mann-Whitney test. (**F**) Experimental schema for analyzing the recall function and phenotype of primary memory cells that developed from shCon and shMll1 transduced P14 cells after initial LCMV_Arm_ infection.

**Fig. S7.**
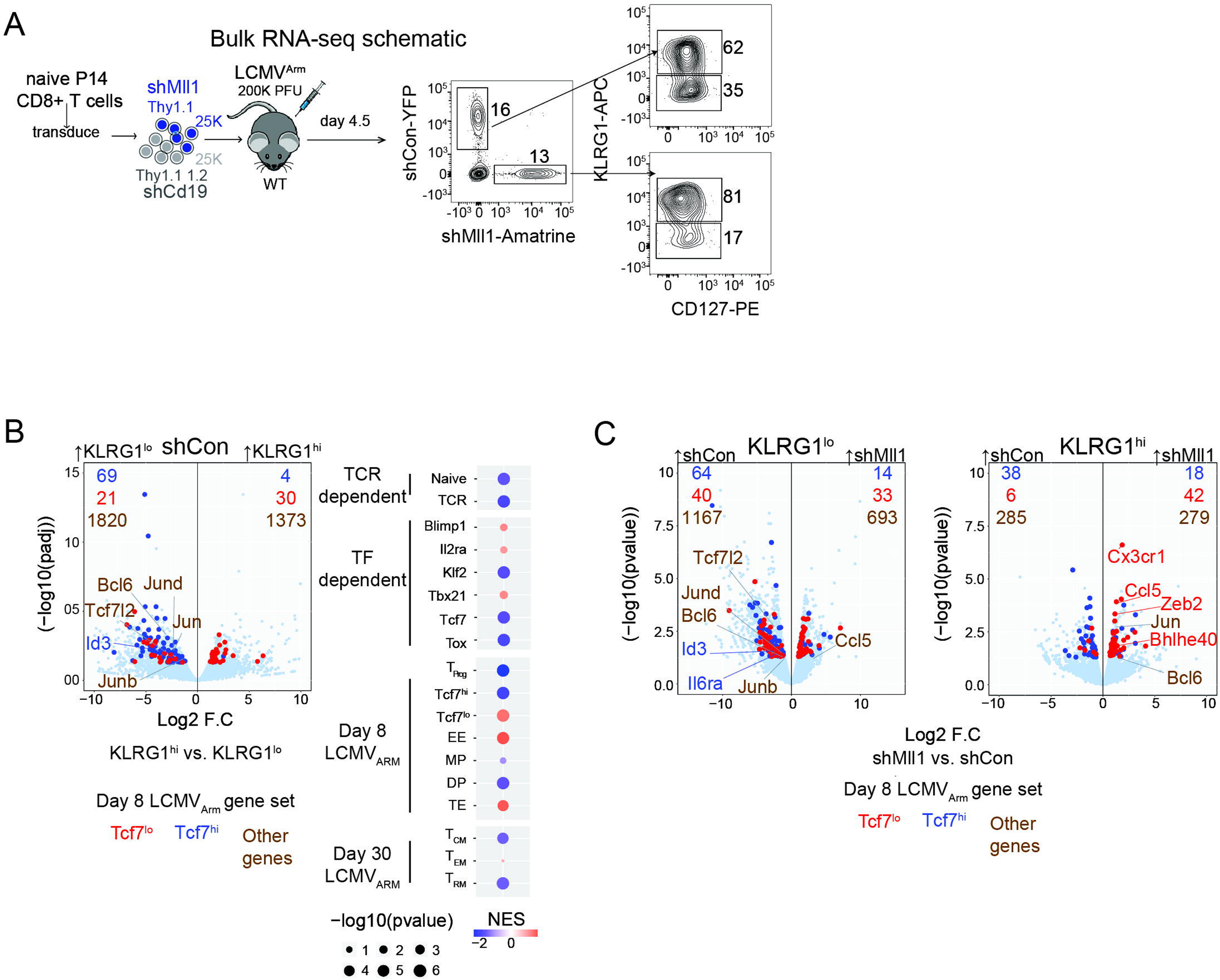
Defective activation of MP and T_SCM_ gene expression by Mll1-deficient P14 cells early after LCMV_Arm_ infection. (**A**) Experimental strategy for bulk RNA-seq of Mll1-deficient P14 cells. (**B**) Transcriptional differences on day 5 pi between wildtype (i.e., shCon-transduced) KLRG1^hi^ and KLRG1^lo^ P14 cells are depicted in the volcano plot. Genes whose expression was significantly different between the KLRG1 subsets that are more highly expressed in Tcf7^hi^ (blue) or Tcf7^lo^ (red) T_EFF_ cells from day 8 after LCMV_Arm_ infection(*12*) were colored and specific signature genes were labeled. Bubble plots depict the relative enrichment of specific gene signatures in gene expression from KLRG1^hi^ vs KLRG1^lo^ cells (Wald statistic). (**C**) Volcano plots show the effect of Mll1 depletion in KLRG1^lo^ (*top*) or KLRG1^hi^ (*bottom*) P14 cells. Genes associated with Tcf7^hi^ (blue) and Tcf7^lo^ (red) cells were colored (*12*), and selected signature genes whose expression was significantly different were labelled.

**Fig. S8.**
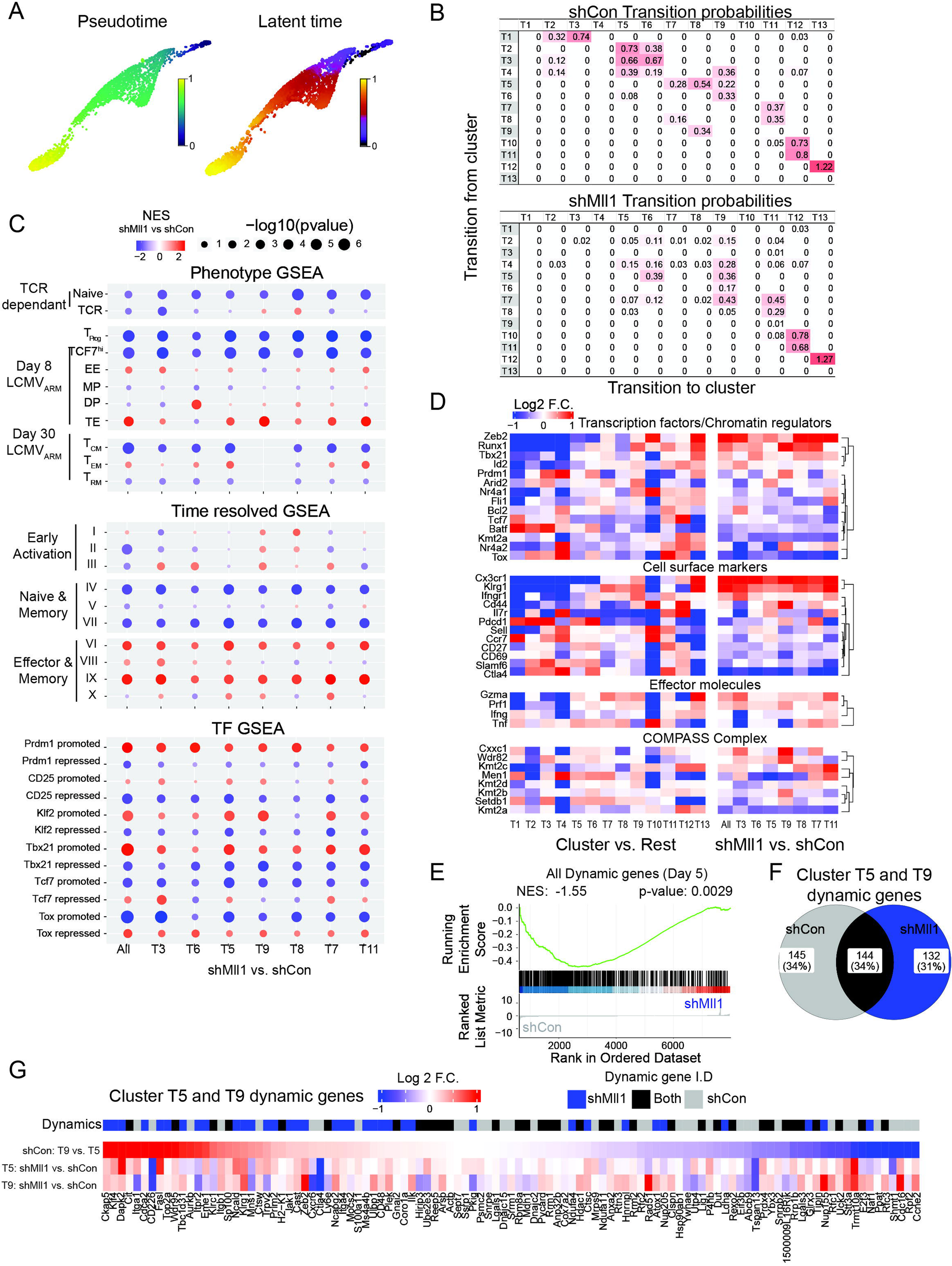
Mll1-deficient P14 cells inefficiently activate TCR-dependent genes whose transcriptional dynamics early during LCMV_Arm_ infection establish development of T_SCM_ precursor cells. **(A)** PAGA analysis of scRNA-seq data overlaid with pseudotime (left) and latent time (right). Pseudotime analysis was performed on all analyzed cells via SCANPY with a random cell in cluster T1 used as the starting node. Latent time was computed on all analyzed cells via scVelo. (B) Tables show cluster specific transition probabilities. Transition probabilities were calculated using data from cells isolated on days 3 and 8 pi, and separately for shCon (top) and shMll1 (bottom) cells that were isolated on day 5 pi with LCMV_Arm_. (C) Bubble plots depict the enrichment of specific gene expression signatures (*left*) (*1, 2, 12, 45, 55, 68, 95*) in gene expression from shCon versus shMll1-transduced cells (Wilcoxon rank score) in each cluster. (D) Heatmap shows the log_2_ fold changes of a subset of T_EFF_ and T_SCM_ signature genes in each Lieden cluster versus all clusters (left), and whose expression differed significantly in shCon and shMll1-transduced cells in the trajectory (right) (Wilcoxon ranked p-value<=0.05). The genes were grouped thematically (labels) and clustered by Euclidean distance in the shMll1 vs. shCon comparison (right). (E) Barcode plot shows the GSEA of all shCon-transduced cell dynamic genes (10^th^ percentile) in the comparison of all shMll1 vs shCon transduced cells. (F) Venn diagram shows overlap of the top dynamic genes (10^th^ percentile) extracted from shCon or shMll1 cells in clusters T5 and T9 that explain their transition probabilities. (G) The presence of the top 10% of dynamic genes identified in either shCon and shMll1 transduced cells from clusters T5 and T9 among (top) all dynamic genes identified in shCon or shMll1 transduced cells from clusters T5 and T9 were determined, and those that were differentially expressed (p<=0.05), their differential expression shMll1 vs shCon transduced cells (bottom) from each cluster was plotted in the heatmap, which was ordered according to their expression between clusters T5 and T9 in shCon-transduced cells (middle).

**Fig. S9.**
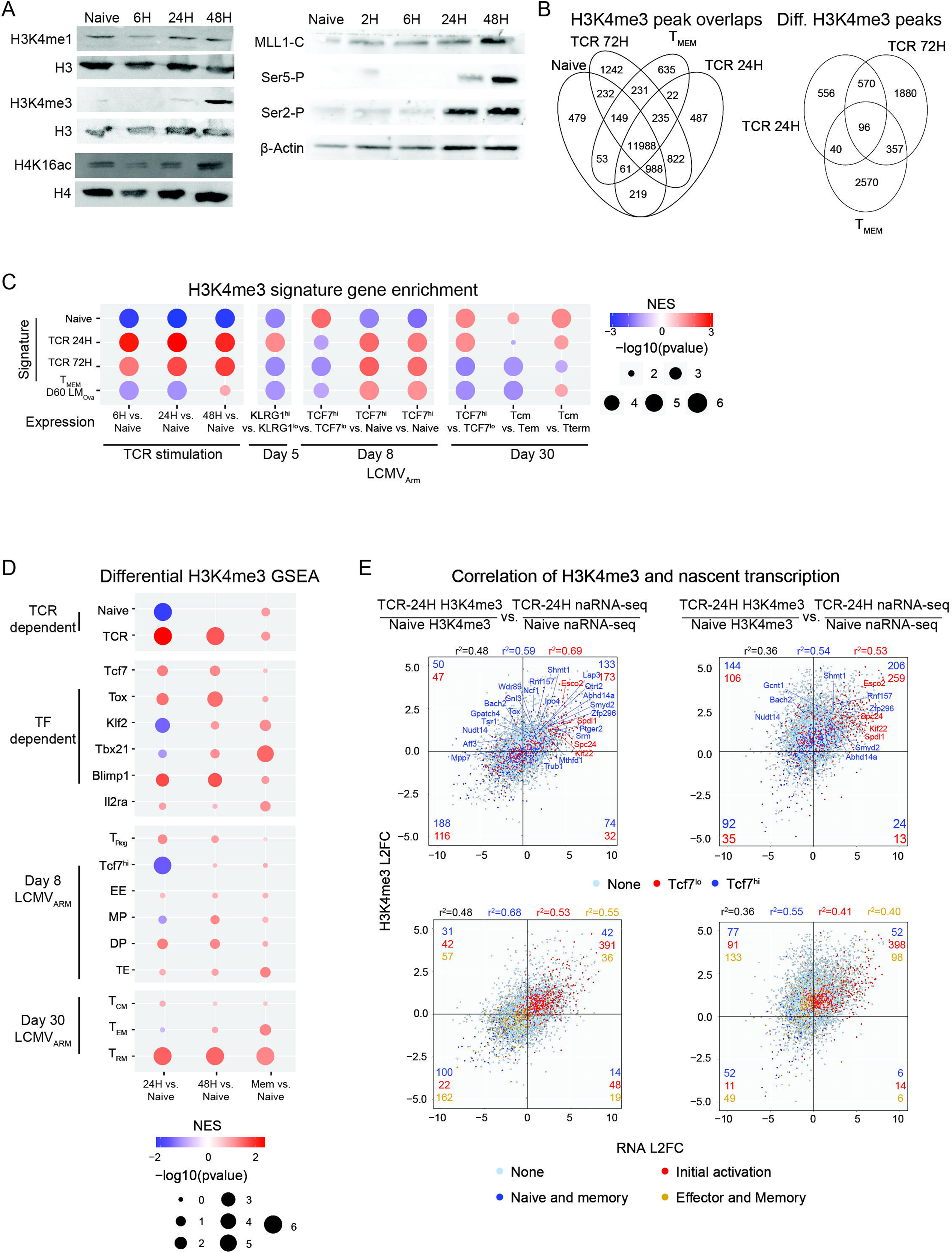
TCR stimulation and Mll1 coordinately establish transcriptional competence by inducing H3K4me3 at transcribed and poised of T_MEM_ cell genes. **(A)** The relative abundance of total and modified histones, Mll1-C and CTD modified RNA Pol II was examined by immunoblotting proteins from the insoluble portion of whole cell RIPA lysates from equivalent numbers of naive cells before and after TCR stimulation. (B) The overlaps between H3K4me3 ChIP-seq peaks in all examined CD8 T cell states (*left*), and specifically for those that exhibit increased H3K4me3 deposition in activated CD8 T cell states compared to naive cells (*right*) are shown in the Venn diagrams. Overlapping peaks had a minimum of one base pair overlap. (C) Bubble plot quantifies enrichment of differential gene expression (Wald statistic) from the indicated comparisons within genes harboring annotated H3K4me3 peaks that exhibited significantly increased or decreased H3K4me3 density in naive cells compared to TCR-activated or T_MEM_ (LM_Ova_ D60) cells (signatures).(D) Bubble plots show the enrichment of genes that exhibit differential H3K4me3 deposition after naive cell activation within specific gene expression signatures (labelled on left). Regions of differential H3K4me3 deposition in activated versus naive cells were annotated to the nearest gene and the average log_2_ fold change was calculated per gene and used for enrichment analysis. (E) The correlation between H3K4me3 deposition (y-axis) and chromatin-associated RNA expression (x-axis) in TCR stimulated versus naive P14 CD8 T cells is shown. Naive cells were stimulated for 24 (*left*) or 72 hours (*right*) before H3K4me3 ChIP-seq and for 24 (*left*) or 48 hours (*right*) before isolation of chromatin-associated RNA. The average log_2_ fold changes in H3K4me3 deposition and chromatin-associated RNA expression were calculated per gene and plotted. Signature genes previously identified as differentially expressed in Tcf7^hi^ (red) or Tcf7^lo^ (blue) T_EFF_ cells 8 days after LCMV_Arm_ infection (*12*)(*top* plots) or involved in initial activation (red), naïve/memory cells (blue), or effector/memory cells (gold), were marked accordingly(*48, 98*) (*bottom* plots), and the number of genes from each category are indicated in each quadrant. R^2^ values for each gene set and for all plotted genes (black) are indicated above each graph.

**Fig. S10.**
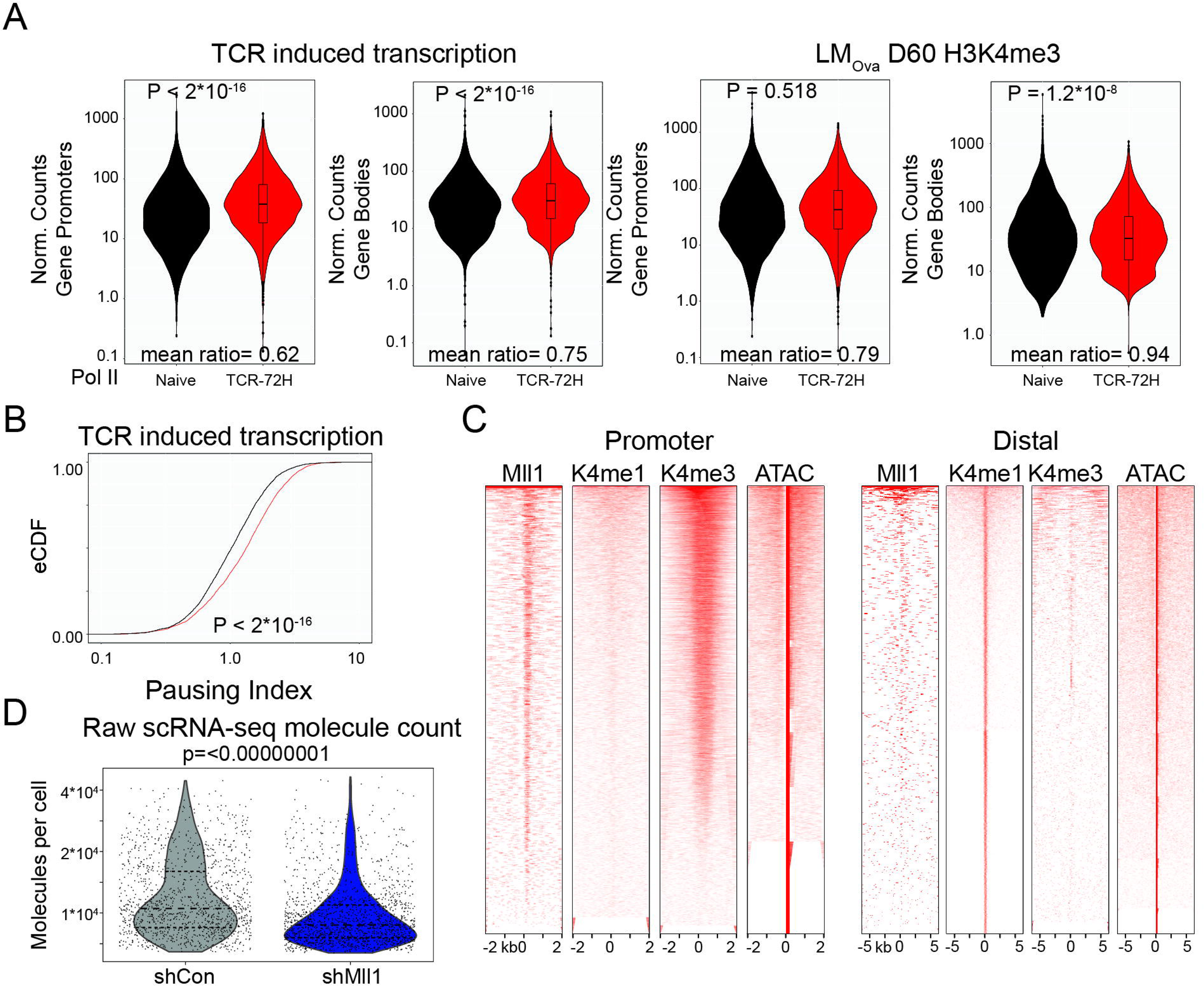
Mll1 preferentially binds active TSSs and promotes transcription of genes *in vivo* that become transcriptionally paused upon TCR stimulation. **(A)** Violin plots show normalized counts of Pol II ChIP-seq reads from naïve (black) or activated (red) cells found in promoters or bodies of genes whose transcription is TCR dependent (*left*) or which have increased H3K4me3 in T_MEM_ relative to naïve cells. (B) The pausing indices of TCR dependent genes from naïve (black) and activated (red) cells are plotted as an eCDF. (C) Heatmaps show relative abundance ChIP-seq reads after mapping Mll1 (ChIP-exo5.0), H3K4me1 and H3K4me3 around accessible promoters and distal enhancers. Promoter regions are TSS’s that are hyperaccessible (ATAC-seq) in activated CD8^+^ T cells (*30*). Distal regions are those that are similarly open, but have a H3K4me1 mark deposited. (D) Violin plots show the number of molecules counted per encapsulated and sequenced cell during scRNA-seq. P value determined via Welch’s unpaired t-test.

## References

1. H. Diao, et al., Single-cell lineage trajectories and chromatin regulators that initialize antiviral CD8 T cell ontogeny. bioRxiv, 2021.2008.2011.456014 (2021).

2. C. Yao et al., Single-cell RNA-seq reveals TOX as a key regulator of CD8+ T cell persistence in chronic infection. Nature Immunology 20, 890-901 (2019).

3. B. Kakaradov et al., Early transcriptional and epigenetic regulation of CD8+ T cell differentiation revealed by single-cell RNA sequencing. Nat Immunol 18, 422–432 (2017).

4. A. C. Richard et al., T cell cytolytic capacity is independent of initial stimulation strength. Nat Immunol 19, 849–858 (2018).

5. J. M. Marchingo, et al., T cell signaling. Antigen affinity, costimulation, and cytokine inputs sum linearly to amplify T cell expansion. Science 346, 1123–1127 (2014).

6. D. Zehn, R. Thimme, E. Lugli, G. P. de Almeida, A. Oxenius, ’Stem-like’ precursors are the fount to sustain persistent CD8(+) T cell responses. Nat Immunol 23, 836–847 (2022).

7. S. C. Jameson, D. Masopust, Understanding Subset Diversity in T Cell Memory. Immunity 48, 214–226 (2018).

8. S. M. Kaech, E. J. Wherry, Heterogeneity and cell-fate decisions in effector and memory CD8+ T cell differentiation during viral infection. Immunity 27, 393–405 (2007).

9. L. M. McLane, M. S. Abdel-Hakeem, E. J. Wherry, CD8 T Cell Exhaustion During Chronic Viral Infection and Cancer. Annu Rev Immunol 37, 457–495 (2019).

10. M. Philip, A. Schietinger, CD8(+) T cell differentiation and dysfunction in cancer. Nat Rev Immunol 22, 209–223 (2022).

11. R. S. Akondy et al., Origin and differentiation of human memory CD8 T cells after vaccination. Nature 552, 362–367 (2017).

12. D. Pais Ferreira et al., Central memory CD8(+) T cells derive from stem-like Tcf7(hi) effector cells in the absence of cytotoxic differentiation. Immunity 53, 985–1000.e1011 (2020).

13. J. B. Johnnidis, et al., Inhibitory signaling sustains a distinct early memory CD8(+) T cell precursor that is resistant to DNA damage. Sci Immunol 6, (2021).

14. P. Graef et al., Serial transfer of single-cell-derived immunocompetence reveals stemness of CD8(+) central memory T cells. Immunity 41, 116–126 (2014).

15. L. Gattinoni et al., A human memory T cell subset with stem cell-like properties. Nat Med 17, 1290–1297 (2011).

16. W. W. Lin et al., CD8(+) T Lymphocyte Self-Renewal during Effector Cell Determination. Cell Rep 17, 1773–1782 (2016).

17. X. Zhou et al., Differentiation and persistence of memory CD8(+) T cells depend on T cell factor 1. Immunity 33, 229–240 (2010).

18. A. G. Soerens et al., Functional T cells are capable of supernumerary cell division and longevity. Nature 614, 762–766 (2023).

19. W. H. Hudson et al., Proliferating Transitory T Cells with an Effector-like Transcriptional Signature Emerge from PD-1(+) Stem-like CD8(+) T Cells during Chronic Infection. Immunity 51, 1043–1058 e1044 (2019).

20. R. Zander et al., CD4(+) T Cell Help Is Required for the Formation of a Cytolytic CD8(+) T Cell Subset that Protects against Chronic Infection and Cancer. Immunity 51, 1028–1042 e1024 (2019).

21. D. T. Utzschneider et al., T Cell Factor 1-Expressing Memory-like CD8+ T Cells Sustain the Immune Response to Chronic Viral Infections. Immunity 45, 415–427 (2016).

22. S. J. Im et al., Defining CD8+ T cells that provide the proliferative burst after PD-1 therapy. Nature 537, 417–421 (2016).

23. R. He et al., Follicular CXCR5-expressing CD8+ T cells curtail chronic viral infection. Nature 537, 412–416 (2016).

24. R. Zander et al., CD4(+) T Cell Help Is Required for the Formation of a Cytolytic CD8(+) T Cell Subset that Protects against Chronic Infection and Cancer. Immunity 51, 1028–1042.e1024 (2019).

25. W. H. Hudson et al., Proliferating Transitory T Cells with an Effector-like Transcriptional Signature Emerge from PD-1+ Stem-like CD8+ T Cells during Chronic Infection. Immunity 51, 1043–1058.e1044 (2019).

26. J.-C. Beltra et al., Developmental Relationships of Four Exhausted CD8+ T Cell Subsets Reveals Underlying Transcriptional and Epigenetic Landscape Control Mechanisms. Immunity 52, 825–841.e828 (2020).

27. B. C. Miller et al., Subsets of exhausted CD8(+) T cells differentially mediate tumor control and respond to checkpoint blockade. Nat Immunol 20, 326–336 (2019).

28. K. E. Pauken et al., Epigenetic stability of exhausted T cells limits durability of reinvigoration by PD-1 blockade. Science 354, 1160–1165 (2016).

29. C. D. Scharer, A. P. Bally, B. Gandham, J. M. Boss, Cutting Edge: Chromatin Accessibility Programs CD8 T Cell Memory. J Immunol 198, 2238–2243 (2017).

30. J. P. Scott-Browne et al., Dynamic Changes in Chromatin Accessibility Occur in CD8(+) T Cells Responding to Viral Infection. Immunity 45, 1327–1340 (2016).

31. B. Yu et al., Epigenetic landscapes reveal transcription factors that regulate CD8(+) T cell differentiation. Nature immunology 18, 573–582 (2017).

32. S. M. Gray, R. A. Amezquita, T. Guan, S. H. Kleinstein, S. M. Kaech, Polycomb Repressive Complex 2-Mediated Chromatin Repression Guides Effector CD8(+) T Cell Terminal Differentiation and Loss of Multipotency. Immunity 46, 596–608 (2017).

33. B. Kakaradov et al., Early transcriptional and epigenetic regulation of CD8(+) T cell differentiation revealed by single-cell RNA sequencing. Nat Immunol 18, 422–432 (2017).

34. L. Pace et al., The epigenetic control of stemness in CD8(+) T cell fate commitment. Science 359, 177–186 (2018).

35. A. Guo et al., cBAF complex components and MYC cooperate early in CD8(+) T cell fate. Nature 607, 135–141 (2022).

36. D. T. Utzschneider et al., Early precursor T cells establish and propagate T cell exhaustion in chronic infection. Nat Immunol 21, 1256–1266 (2020).

37. E. Ahn et al., Role of PD-1 during effector CD8 T cell differentiation. Proc Natl Acad Sci U S A 115, 4749–4754 (2018).

38. M. E. Pipkin et al., Interleukin-2 and inflammation induce distinct transcriptional programs that promote the differentiation of effector cytolytic T cells. Immunity 32, 79–90 (2010).

39. B. K. Cenik, A. Shilatifard, COMPASS and SWI/SNF complexes in development and disease. Nat Rev Genet 22, 38–58 (2021).

40. C. M. Hughes et al., Menin associates with a trithorax family histone methyltransferase complex and with the hoxc8 locus. Mol Cell 13, 587–597 (2004).

41. A. Yokoyama et al., Leukemia proto-oncoprotein MLL forms a SET1-like histone methyltransferase complex with menin to regulate Hox gene expression. Mol Cell Biol 24, 5639–5649 (2004).

42. J. Paggetti et al., Crosstalk between leukemia-associated proteins MOZ and MLL regulates HOX gene expression in human cord blood CD34+ cells. Oncogene 29, 5019–5031 (2010).

43. S. Denissov et al., Mll2 is required for H3K4 trimethylation on bivalent promoters in embryonic stem cells, whereas Mll1 is redundant. Development 141, 526–537 (2014).

44. Y. Ji et al., Repression of the DNA-binding inhibitor Id3 by Blimp-1 limits the formation of memory CD8+ T cells. Nature Immunology 12, 1230–1237 (2011).

45. J. A. Best et al., Transcriptional insights into the CD8(+) T cell response to infection and memory T cell formation. Nat Immunol 14, 404–412 (2013).

46. E. J. Wherry, J. N. Blattman, K. Murali-Krishna, R. van der Most, R. Ahmed, Viral persistence alters CD8 T-cell immunodominance and tissue distribution and results in distinct stages of functional impairment. J Virol 77, 4911–4927 (2003).

47. S. M. Kaech, S. Hemby, E. Kersh, R. Ahmed, Molecular and functional profiling of memory CD8 T cell differentiation. Cell 111, 837–851 (2002).

48. C. Y. Yang et al., The transcriptional regulators Id2 and Id3 control the formation of distinct memory CD8+ T cell subsets. Nat Immunol 12, 1221–1229 (2011).

49. W.-Hsuan W. Lin, et al., CD8+ T Lymphocyte Self-Renewal during Effector Cell Determination. Cell Reports 17, 1773–1782 (2016).

50. V. Kalia et al., Prolonged interleukin-2Ralpha expression on virus-specific CD8+ T cells favors terminal-effector differentiation in vivo. Immunity 32, 91–103 (2010).

51. N. S. Joshi et al., Inflammation directs memory precursor and short-lived effector CD8(+) T cell fates via the graded expression of T-bet transcription factor. Immunity 27, 281–295 (2007).

52. C. Gerlach et al., The chemokine receptor CX3CR1 defines three antigen-experienced CD8 T cell subsets with distinct roles in immune surveillance and homeostasis. Immunity 45, 1270–1284 (2016).

53. J. J. Obar, K. M. Khanna, L. Lefrançois, Endogenous naive CD8+ T cell precursor frequency regulates primary and memory responses to infection. Immunity 28, 859–869 (2008).

54. E. J. Wherry et al., Lineage relationship and protective immunity of memory CD8 T cell subsets. Nature immunology 4, 225–234 (2003).

55. D. Wang et al., The Transcription Factor Runx3 Establishes Chromatin Accessibility of <em>cis</em>-Regulatory Landscapes that Drive Memory Cytotoxic T Lymphocyte Formation. Immunity 48, 659-674.e656 (2018).

56. V. Bergen, M. Lange, S. Peidli, F. A. Wolf, F. J. Theis, Generalizing RNA velocity to transient cell states through dynamical modeling. Nature Biotechnology 38, 1408–1414 (2020).

57. M. G. Guenther, S. S. Levine, L. A. Boyer, R. Jaenisch, R. A. Young, A Chromatin Landmark and Transcription Initiation at Most Promoters in Human Cells. Cell 130, 77–88 (2007).

58. H. Wang et al., H3K4me3 regulates RNA polymerase II promoter-proximal pause-release. Nature 615, 339–348 (2023).

59. B. E. Russ et al., Distinct epigenetic signatures delineate transcriptional programs during virus-specific CD8(+) T cell differentiation. Immunity 41, 853–865 (2014).

60. T. A. Milne et al., MLL Targets SET Domain Methyltransferase Activity to Hox Gene Promoters. Molecular Cell 10, 1107–1117 (2002).

61. T. Nakamura et al., ALL-1 Is a Histone Methyltransferase that Assembles a Supercomplex of Proteins Involved in Transcriptional Regulation. Molecular Cell 10, 1119–1128 (2002).

62. P. Wang et al., Global analysis of H3K4 methylation defines MLL family member targets and points to a role for MLL1-mediated H3K4 methylation in the regulation of transcriptional initiation by RNA polymerase II. Mol Cell Biol 29, 6074–6085 (2009).

63. M. Wu et al., Molecular regulation of H3K4 trimethylation by Wdr82, a component of human Set1/COMPASS. Mol Cell Biol 28, 7337–7344 (2008).

64. J. J. Milner et al., Delineation of a molecularly distinct terminally differentiated memory CD8 T cell population. Proc Natl Acad Sci U S A 117, 25667–25678 (2020).

65. M. J. Rossi, W. K. M. Lai, B. F. Pugh, Simplified ChIP-exo assays. Nat Commun 9, 2842 (2018).

66. E.-J. Cho, T. Takagi, C. R. Moore, S. Buratowski, mRNA capping enzyme is recruited to the transcription complex by phosphorylation of the RNA polymerase II carboxy-terminal domain. Genes & development 11, 3319–3326 (1997).

67. H. H. Ng, F. Robert, R. A. Young, K. Struhl, Targeted recruitment of Set1 histone methylase by elongating Pol II provides a localized mark and memory of recent transcriptional activity. Mol Cell 11, 709–719 (2003).

68. L. K. Mackay et al., Hobit and Blimp1 instruct a universal transcriptional program of tissue residency in lymphocytes. Science 352, 459–463 (2016).

69. R. Chen et al., In vivo RNA interference screens identify regulators of antiviral CD4(+) and CD8(+) T cell differentiation. Immunity 41, 325–338 (2014).

70. C. Fellmann et al., An Optimized microRNA Backbone for Effective Single-Copy RNAi. Cell Reports 5, 1704–1713 (2013).

71. Simon R. V. Knott et al., A Computational Algorithm to Predict shRNA Potency. Molecular Cell 56, 796–807.

72. K. Labun et al., CHOPCHOP v3: expanding the CRISPR web toolbox beyond genome editing. Nucleic Acids Research 47, W171–W174 (2019).

73. Q. Yang et al., TCF-1 upregulation identifies early innate lymphoid progenitors in the bone marrow. Nature Immunology 16, 1044–1050 (2015).

74. H. Pircher, K. Burki, R. Lang, H. Hengartner, R. M. Zinkernagel, Tolerance induction in double specific T-cell receptor transgenic mice varies with antigen. Nature 342, 559–561 (1989).

75. P. Mombaerts et al., Mutations in T-cell antigen receptor genes α and β block thymocyte development at different stages. Nature 360, 225–231 (1992).

76. clusterProfiler: an R Package for Comparing Biological Themes Among Gene Clusters. OMICS: A Journal of Integrative Biology 16, 284-287 (2012).

77. R. Ahmed, A. Salmi, L. D. Butler, J. M. Chiller, M. B. Oldstone, Selection of genetic variants of lymphocytic choriomeningitis virus in spleens of persistently infected mice. Role in suppression of cytotoxic T lymphocyte response and viral persistence. J Exp Med 160, 521–540 (1984).

78. J. A. Olson, C. McDonald-Hyman, S. C. Jameson, S. E. Hamilton, Effector-like CD8(+) T cells in the memory population mediate potent protective immunity. Immunity 38, 1250–1260 (2013).

79. J. J. Milner et al., Bromodomain protein BRD4 directs and sustains CD8 T cell differentiation during infection. J Exp Med 218, (2021).

80. J. J. Milner et al., Runx3 programs CD8+ T cell residency in non-lymphoid tissues and tumours. Nature 552, 253 (2017).

81. L. B. Rodda et al., Single-Cell RNA Sequencing of Lymph Node Stromal Cells Reveals Niche-Associated Heterogeneity. Immunity 48, 1014–1028.e1016 (2018).

82. R. Patro, G. Duggal, M. I. Love, R. A. Irizarry, C. Kingsford, Salmon: fast and bias-aware quantification of transcript expression using dual-phase inference. Nature methods 14, 417–419 (2017).

83. M. I. Love, W. Huber, S. Anders, Moderated estimation of fold change and dispersion for RNA-seq data with DESeq2. Genome Biol 15, 550 (2014).

84. C. Yao et al., Single-cell RNA-seq reveals TOX as a key regulator of CD8(+) T cell persistence in chronic infection. Nat Immunol 20, 890–901 (2019).

85. F. A. Wolf, P. Angerer, F. J. Theis, SCANPY: large-scale single-cell gene expression data analysis. Genome Biology 19, 15 (2018).

86. L. Haghverdi, A. T. L. Lun, M. D. Morgan, J. C. Marioni, Batch effects in single-cell RNA-sequencing data are corrected by matching mutual nearest neighbors. Nature Biotechnology 36, 421–427 (2018).

87. F. A. Wolf et al., PAGA: graph abstraction reconciles clustering with trajectory inference through a topology preserving map of single cells. Genome Biology 20, 59 (2019).

88. M. J. Rossi, W. K. M. Lai, B. F. Pugh, Simplified ChIP-exo assays. Nature Communications 9, 2842 (2018).

89. B. Langmead, S. L. Salzberg, Fast gapped-read alignment with Bowtie 2. Nat Methods 9, 357–359 (2012).

90. P. Danecek et al., Twelve years of SAMtools and BCFtools. Gigascience 10, (2021).

91. Y. Zhang et al., Model-based Analysis of ChIP-Seq (MACS). Genome Biology 9, R137 (2008).

92. C. S. Ross-Innes et al., Differential oestrogen receptor binding is associated with clinical outcome in breast cancer. Nature 481, 389–393 (2012).

93. G. Yu, L.-G. Wang, Q.-Y. He, ChIPseeker: an R/Bioconductor package for ChIP peak annotation, comparison and visualization. Bioinformatics 31, 2382–2383 (2015).

94. C. X. Dominguez et al., The transcription factors ZEB2 and T-bet cooperate to program cytotoxic T cell terminal differentiation in response to LCMV viral infection. J Exp Med 212, 2041–2056 (2015).

95. A. Xin et al., A molecular threshold for effector CD8(+) T cell differentiation controlled by transcription factors Blimp-1 and T-bet. Nat Immunol 17, 422–432 (2016).

96. Z. Gu, R. Eils, M. Schlesner, Complex heatmaps reveal patterns and correlations in multidimensional genomic data. Bioinformatics 32, 2847–2849 (2016).

97. P. B. Rahl et al., c-Myc regulates transcriptional pause release. Cell 141, 432–445 (2010).

98. H. Diao, M. Pipkin, Stability and flexibility in chromatin structure and transcription underlies memory CD8 T-cell differentiation [version 1; peer review: 2 approved]. F1000Research 8, (2019).

